# Development of a clostridia-based cell-free system for prototyping genetic parts and metabolic pathways

**DOI:** 10.1101/2020.03.11.987826

**Authors:** Antje Krüger, Alexander P. Mueller, Grant A. Rybnicky, Nancy L. Engle, Zamin K. Yang, Tim J. Tschaplinski, Sean D. Simpson, Michael Köpke, Michael C. Jewett

## Abstract

Gas fermentation by autotrophic bacteria, such as clostridia, offers a sustainable path to numerous bioproducts from a range of local, highly abundant, waste and low-cost feedstocks, such as industrial flue gases or syngas generated from biomass or municipal waste. Unfortunately, designing and engineering clostridia remains laborious and slow. The ability to prototype individual genetic parts, gene expression, and biosynthetic pathway performance *in vitro* before implementing them in cells could help address these bottlenecks by speeding up design. Unfortunately, a high-yielding cell-free gene expression (CFE) system from clostridia has yet to be developed. Here, we report the development and optimization of a high-yielding (236 ± 24 µg/mL) batch CFE platform from the industrially relevant anaerobe, *Clostridium autoethanogenum.* A key feature of the platform is that both circular and linear DNA templates can be applied directly to the CFE reaction to program protein synthesis. We demonstrate the ability to prototype gene expression, and quantitatively map cell-free metabolism in lysates from this system. We anticipate that the *C. autoethanogenum* CFE platform will not only expand the protein synthesis toolkit for synthetic biology, but also serve as a platform in expediting the screening and prototyping of gene regulatory elements in non-model, industrially relevant microbes.

## INTRODUCTION

Microbes can be engineered to manufacture biofuels and high-value compounds such as chemicals, materials, and therapeutics (Keasling, 2012; Nielsen and Keasling, 2016). This biomanufacturing capability promises to help address rapid population growth, an increase in energy demand, and waste generation (Nielsen et al., 2014). However, even the most advanced design-build-test cycles for optimizing a given compound’s biosynthetic pathway in model organisms such as *Escherichia coli* and yeast are still on the order of weeks to months. In addition, process-based challenges associated with these organisms remain (e.g., limited substrate range, reduced yields through CO_2_ losses, and susceptibility to contamination, among others) (Keasling, 2012; Nielsen and Keasling, 2016). These challenges have prevented a more rapid commercialization of new bioproduct manufacturing processes, with only a handful successfully commercialized to date apart from ethanol fermentation (Meadows et al., 2016; Nakamura and Whited, 2003; Nielsen et al., 2014; Yim et al., 2011). As such, most industrial bioprocesses (e.g., synthesis of amino acids (Leuchtenberger et al., 2005), acetone-butanol-ethanol (ABE) (Jiang et al., 2015; Jones, 2005), organic acids (Ghaffar et al., 2014; Rodriguez et al., 2014; Wee et al., 2006)) rely on other “non-model” organisms.

Clostridia are one such group of organisms, which are industrially proven and have exceptional substrate and metabolite diversity, as well as tolerance to metabolic end-products and contaminants (Tracy et al., 2012). Industrial, large-scale fermentations with clostridia have been carried out for over 100 years with the ABE fermentation being the second largest industrial fermentation process only behind ethanol fermentation (Jones, 2005). In addition to ABE clostridia (solventogenic), there are also clostridia species that are able to degrade lignocellulosic biomass (cellulolytic) and species that are capable of autotrophic growth on C1 substrates, such as carbon monoxide (CO) and CO_2_ (acetogenic) (Tracy et al., 2012). Gas fermentation with acetogenic clostridia offers an attractive route for conversion of syngas that can be generated from any biomass resource (e.g., agricultural waste or unsorted and non-recyclable municipal solid waste) and industrial waste resources (e.g., off-gases from steel mills, processing plants or refineries) to fuels and chemicals. However, the current state-of-the-art strain engineering for clostridia remains a low-throughput, labor-intensive endeavor. Specific challenges include organism-specific genetic constraints (Daniell et al., 2015; Joseph et al., 2018; Liew et al., 2017, 2016; Nagaraju et al., 2016), the requirement of an anaerobic environment, and, in case of acetogens, handling of gases. As a result, developments in clostridia biotechnology and basic knowledge of clostridia biology have lagged behind achievements in aerobic prokaryotic and eukaryotic biology. New robust tools are needed to study clostridia and speed up the designing, building, and testing of biological processes in these organisms.

Extract-based cell-free systems are emerging as powerful platforms for synthetic biology applications such as metabolic engineering (Bujara et al., 2011; Carlson et al., 2012; Dudley et al., 2019; Hodgman and Jewett, 2012; Karim et al., 2019a; Karim and Jewett, 2016; Kelwick et al., 2017; Morgado et al., 2018; Silverman et al., 2019a). Assembling metabolic pathways in the cell-free environment has been done traditionally by assembling purified enzymes and substrates. However, the development of cell-free gene expression (CFE) systems has transformed the way pathways can be built and tested (Silverman et al., 2019a). These systems consist of crude cell extracts, energy substrates, co-factors and genetic instructions in the form of DNA, and facilitate the activation, manipulation and usage of cellular processes in a test tube. While cell-free systems have historically been used to study fundamental biology (e.g., the genetic code) (Nirenberg and Matthaei, 1961), recent development of cell-free protein synthesis capabilities (Caschera and Noireaux, 2014; Des Soye et al., 2019; Jewett et al., 2008; Jewett and Swartz, 2004) has expanded the application space to include prototyping of genetic parts (Chappell et al., 2013; Moore et al., 2018; Siegal-Gaskins et al., 2014; Melissa K Takahashi et al., 2015; Melissa K. Takahashi et al., 2015; Yim et al., 2019) and studying whole metabolic pathways (Bujara et al., 2011; Dudley et al., 2019; Karim et al., 2019a; Karim and Jewett, 2016; Kelwick et al., 2017). As compared to *in vivo* approaches, cell-free systems have several key advantages: First, these systems lack a cell wall, and thereby allow active monitoring, rapid sampling and direct manipulation. Second, because genetic instructions can be simply added to CFE reactions in form of plasmid DNA or linear PCR products, they circumvent laborious cloning and transformation steps, and can thereby facilitate testing of genetic designs within a few hours instead of several days or weeks. Third, this approach does not rely on time-consuming enzyme purification procedures but rapidly builds and tests metabolic pathways directly in cell extracts by synthesizing required enzymes *in vitro* (Karim et al., 2018; Karim and Jewett, 2016). Given these advantages, cell-free systems have emerged as an important approach for accelerating biological design, especially with the advent of new extract based systems from non-model organisms: *Bacillus* (Moore et al., 2018), *Streptomyces* (Li et al., 2018, 2017), *Vibrio* (Des Soye et al., 2018; Failmezger et al., 2018; Wiegand et al., 2018), and *Pseudomonas* (Wang et al., 2018) among others. However, no clostridia cell-free system exists that produces protein yields sufficient for prototyping genetic parts and metabolic pathways.

Here, we present the first, to our knowledge, easy-to-use, robust and high-yielding clostridia CFE platform derived from an industrially relevant strain, *Clostridium autoethanogenum* (*C. auto*), which promises to facilitate metabolic engineering applications. Specifically, the goal was to enable cell-free protein synthesis yields of more than 100 μg/ml by systematically optimizing process parameters. To achieve this goal, we first streamline and optimize the extract preparation and processing procedure. Second, we carried out a systematic optimization of CFE reaction conditions to tune the physicochemical environment for stimulating highly active combined transcription and translation from linear DNA templates. We observed a ∼100,000-fold increase in protein synthesis yields relative to the original unoptimized case, resulting in a final titer of approximately 240 μg/mL in 3-h batch reaction. This yield could be further improved in a semi-continuous reaction to more than 300 μg/mL. Finally, with the clostridia CFE system at hand, we demonstrate the capability of our system for clostridia-specific prototyping: clostridia genetic parts by expressing luciferase from constructs under the control of endogenous promoters and 5’UTRs derived from clostridia metabolic enzymes or by utilizing different gene coding sequences, as well as activity of clostridia metabolic pathways in the extracts (**Figure 1**). We anticipate that this platform, the first high-yielding CFE system from an obligate anaerobe, will speed-up metabolic engineering efforts for bioprocess development in clostridia.

**Figure 1.**
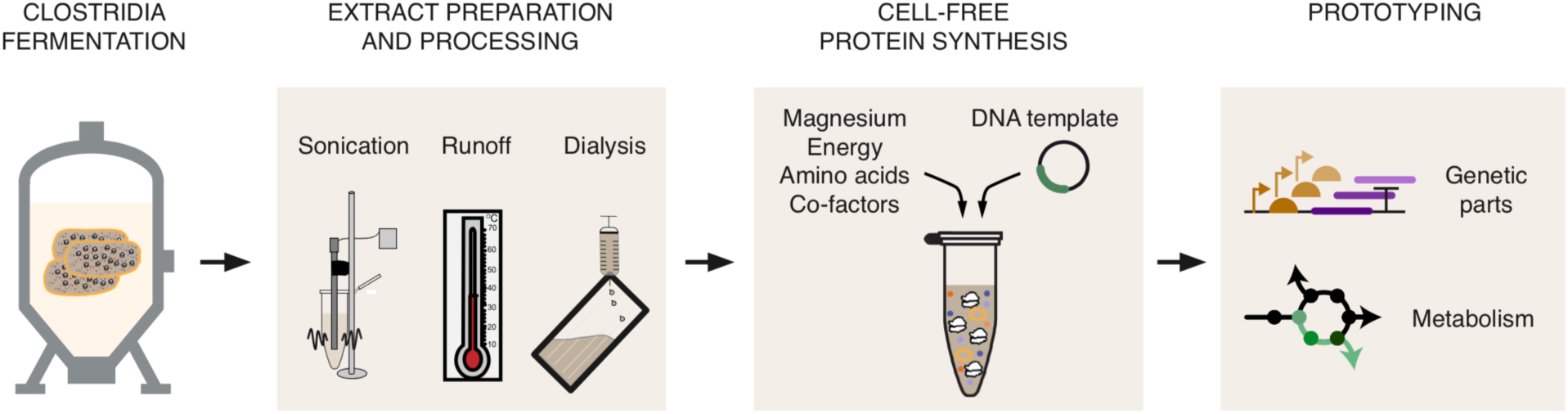
Development of a simple, robust and high-yielding clostridia cell-free gene expression (CFE) platform. Schematic illustration of the development strategy. First, starting from cell pellets collected from clostridia cultures, we initially optimized extract preparation and processing by testing different sonication, runoff and dialysis conditions. Second, we adjusted concentrations of key components in the CFE reaction to further maximize protein production. Third, this optimized CFE system was used to prototype clostridia genetic parts and assess cell-free metabolism.

## RESULTS

Developing a system capable of CFE from a new organism requires optimization at several levels. The choice of organism, fermentation conditions, extract preparation and processing, and cell-free reaction conditions each play an important role. In this work, we aimed to develop a high-yielding CFE system using an industrially relevant clostridia strain as our source organism, *C. auto*. Based on extensive optimization that has gone into establishing anaerobic fermentation conditions for this organism (Heijstra et al., 2017; Valgepea et al., 2017), we chose to fix microbial growth and harvest conditions. Below, we describe our efforts to (i) establish the clostridia-based CFE system, (ii) identify beneficial extract processing steps, and (iii) optimize reaction conditions to enable prototyping of clostridia-based genetic parts and metabolism in the cell-free environment.

### CFE using C. auto extracts requires high Mg(Glu)_2_ concentrations

We started development of *C. auto*-based cell-free systems by exploring the CFE capability when prepared under aerobic conditions and using extract preparation and gene expression conditions of the high-yielding BL21 *E. coli* system (Kwon and Jewett, 2015). In brief, we resuspended *C. auto* cells in buffer containing acetate salts, lysed them by sonication using 640 J total sonication input energy per mL cell suspension, and centrifuged them at 12,000 x g to clarify the lysate (**Figure 2A, left panel**). The resulting extract was used for CFE at 30 °C driven by the PANOx-SP energy regeneration system (Jewett and Swartz, 2004) and containing 8 mM Mg(Glu)_2_, 33 mM phosphoenolpyruvate (PEP), 2 mM of all cognate amino acids, 0.33 mM reduced nicotinamide adenine dinucleotide (NAD^+^) and 0.27 mM coenzyme A (CoA) (**Figure 2A, middle**). We chose firefly luciferase as reporter protein, as it has been demonstrated in clostridia (Feustel et al., 2004) and its expression can be detected via a highly sensitive bioluminescence assay. For this, we cloned a clostridia-codon-adapted variant of the firefly luciferase gene into our CFE expression vector pJL1 under control of the T7 promoter, added the construct to the CFE reaction, and followed luciferase expression in CFE by luminescence for almost 3 hours. We observed little to no luminescence above a no plasmid control (**Figure 2A, right panel**).

Given the poor protein synthesis yields, we next performed a magnesium optimization as it has been shown to be one of the most critical factors impacting CFE productivity (Des Soye et al., 2018; Hodgman and Jewett, 2013; Jewett and Swartz, 2004; Kwon and Jewett, 2015; Li et al., 2017; Martin et al., 2017; Wang et al., 2018). We set up CFE reactions over a range of Mg(Glu)_2_ concentrations between 8 mM and 36 mM. Mg(Glu)_2_ concentrations of ≥ 20 mM markedly increased luciferase expression by more than five orders of magnitude (**Figure 2B**) with the optimum at 32 mM Mg(Glu)_2_. This result was surprising because the optimum for *E. coli* extracts tends to be in the range of 8 mM to 12 mM Mg(Glu)_2_ (Jewett and Swartz, 2004; Kwon and Jewett, 2015). While we do not understand the requirement for high magnesium concentrations for protein synthesis, we subsequently carried out a series of optimization experiments to explore a range of both process and reaction conditions to improve cell-free performance.

**Figure 2.**
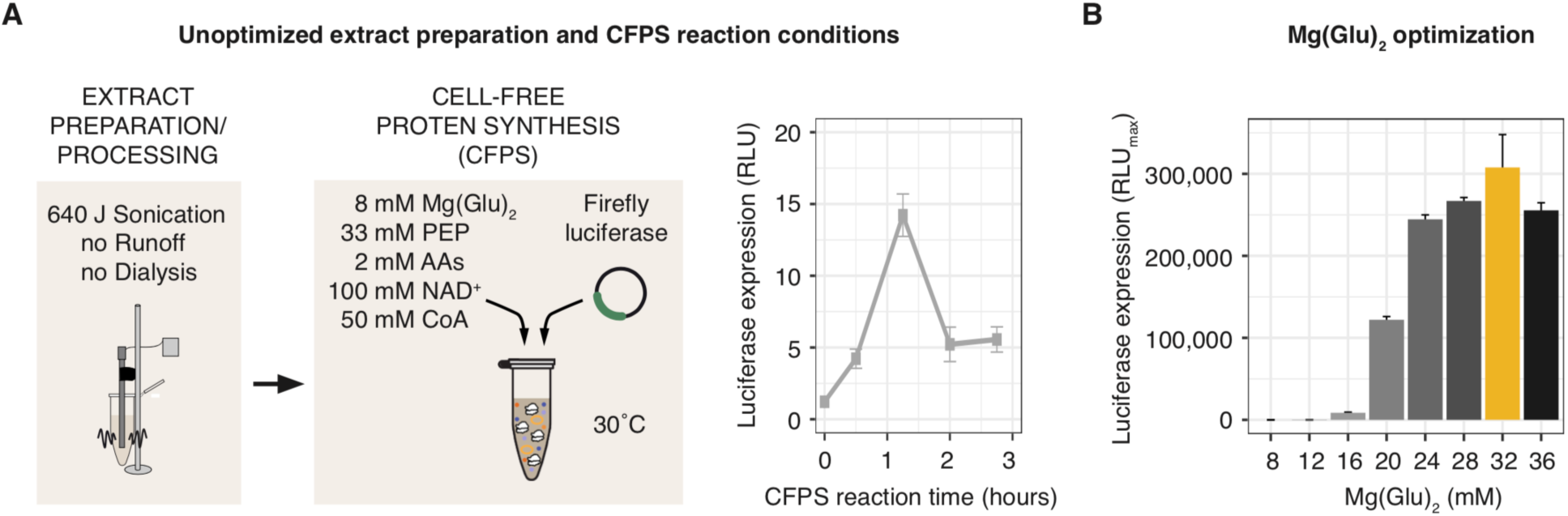
*C. auto*-derived CFE requires different conditions than *E. coli*-derived CFE. (A) Using *E. coli* conditions for extract preparation/ processing and CFE reactions, luciferase expression was determined in *C. auto* extracts. (A, left and middle panel) Simplified schematic of extract preparation and processing steps and key components of CFE reactions, respectively. (A, right panel) Luciferase expression in *C. auto* extracts during CFE at 8 mM Mg(Glu)_2_. (B) Maximum luciferase expression during CFE at different Mg(Glu)_2_ concentrations. Yellow bar indicates optimized condition. *C. auto* cell pellets were resuspended in S30 buffer, lysed by sonication at 640 J, clarified by centrifugation at 12,000 x g, and used for CFE containing the key components at indicated concentrations in (A). Luciferase expression was determined by bioluminescence. PEP: phosphoenolpyruvate; AAs: amino acids; NAD^+^: reduced nicotinamide adenine dinucleotide; CoA: coenzyme A. Data are presented as mean ± s.d. of at least three independent reactions.

### Adjusting extract preparation and processing of C. auto increases CFE yields

The quality of prepared crude cell extract, which is largely determined by how the cells are lysed and processed (*i.e.,* run-off reactions, dialysis), has a significant effect on CFE (Carlson et al., 2012; Gregorio et al., 2019; Kim et al., 1996; Kwon and Jewett, 2015; Silverman et al., 2019b). We therefore explored key parameters of both (**Figure 3A**), starting with lysis conditions responsible for cell wall rupture. Using sonication as our lysis method due to its simple, reproducible, and inexpensive nature (Kwon and Jewett, 2015), we lysed 1 mL of resuspended *C. auto* cells at different sonication input energies ranging from 250 J to 910 J at 50% amplitude for 10 sec on and 10 sec off (**Figure 3B**). We clarified the lysates by centrifugation and tested the extract’s capability for CFE. Compared to the initially used 640 J, higher input energies reduced CFE yields, while lower energies were beneficial. We found the optimum to be 350 J, which increased luciferase expression by ∼30%.

**Figure 3.**
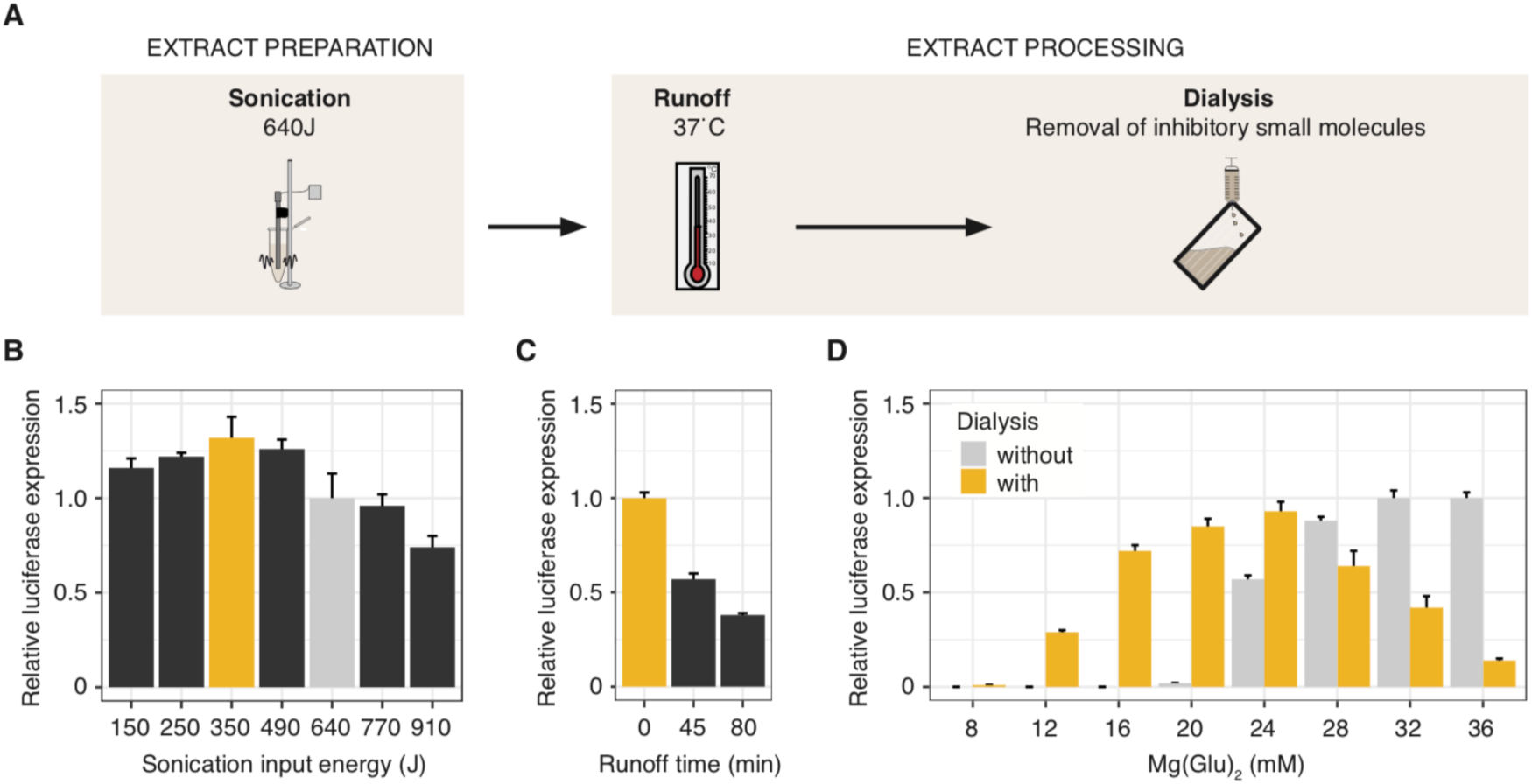
Optimization of *C. auto* extract preparation and processing. (A) Schematic diagram of the extract preparation workflow. (B-D) Relative maximum luciferase luminescence *in vitro* from *C. auto* extracts during 4.25 h CFE prepared using (B) indicated sonication input energies, (C) 350 J sonication input energy and indicated runoff times, and (D) 350J sonication input energy without and with dialysis. Light grey and yellow bars indicate previously and newly optimized condition, respectively. Luciferase expression was determined by bioluminescence and plotted as relative values compared to the maximal luciferase expression of the previously used condition. Data are presented as mean ± s.d. of three independent reactions.

Two common post-lysis processing steps, runoff and dialysis, can improve the quality of extracts for CFE. The runoff involves incubating the extract at 37 °C, which can increase the extract’s protein synthesis productivity (Kwon and Jewett, 2015). The extra time at a physiological temperature is hypothesized to allow ribosomes to “run off” native mRNAs, which might then be degraded by endogenous RNases while the ribosomes are freed-up for synthesis of recombinant proteins (Jermutus et al., 1998; Nirenberg and Matthaei, 1961). To test the effect of a runoff step, we incubated the clarified lysates after sonication at 37 °C for a short (45 min) and a long (80 min) time, clarified them a second time by centrifugation at 12,000 x g and compared their protein synthesis activity. We found that the runoff markedly decreased luciferase expression (**Figure 3C**). A runoff for 45 min almost halved luciferase amounts as compared to no runoff, while longer incubation time (80 min) reduced yields to a third.

In contrast to runoff, dialysis changes the extract’s composition by allowing exchange of small molecules between a dialysis buffer and the extract. This step can be beneficial by removing small molecule inhibitors to increase CFE yields (Gregorio et al., 2019; Silverman et al., 2019b). To test the impact of dialysis, we dialyzed the clarified lysates after sonication three times for 45 min each in S30 buffer at 4 °C and compared luciferase expression at several Mg(Glu)_2_ concentrations. We found that dialysis did not significantly affect overall extract productivity but instead decreased the Mg(Glu)_2_ optimum from 32 mM to 24 mM (**Figure 3D**). Based on these results, we next set out to optimize CFE reaction conditions with an extract preparation and processing protocol that now includes dialysis.

### Adapting CFE reaction conditions further improved C. auto extract-based CFE

As a means to further increase CFE yields, we next optimized several well-known cell-free reaction parameters including the energy regeneration system, the amino acid concentration, co-factor concentration, the extract concentration and oxygen availability, the DNA template, and reaction temperature (**Figure 4A**).

**Figure 4:**
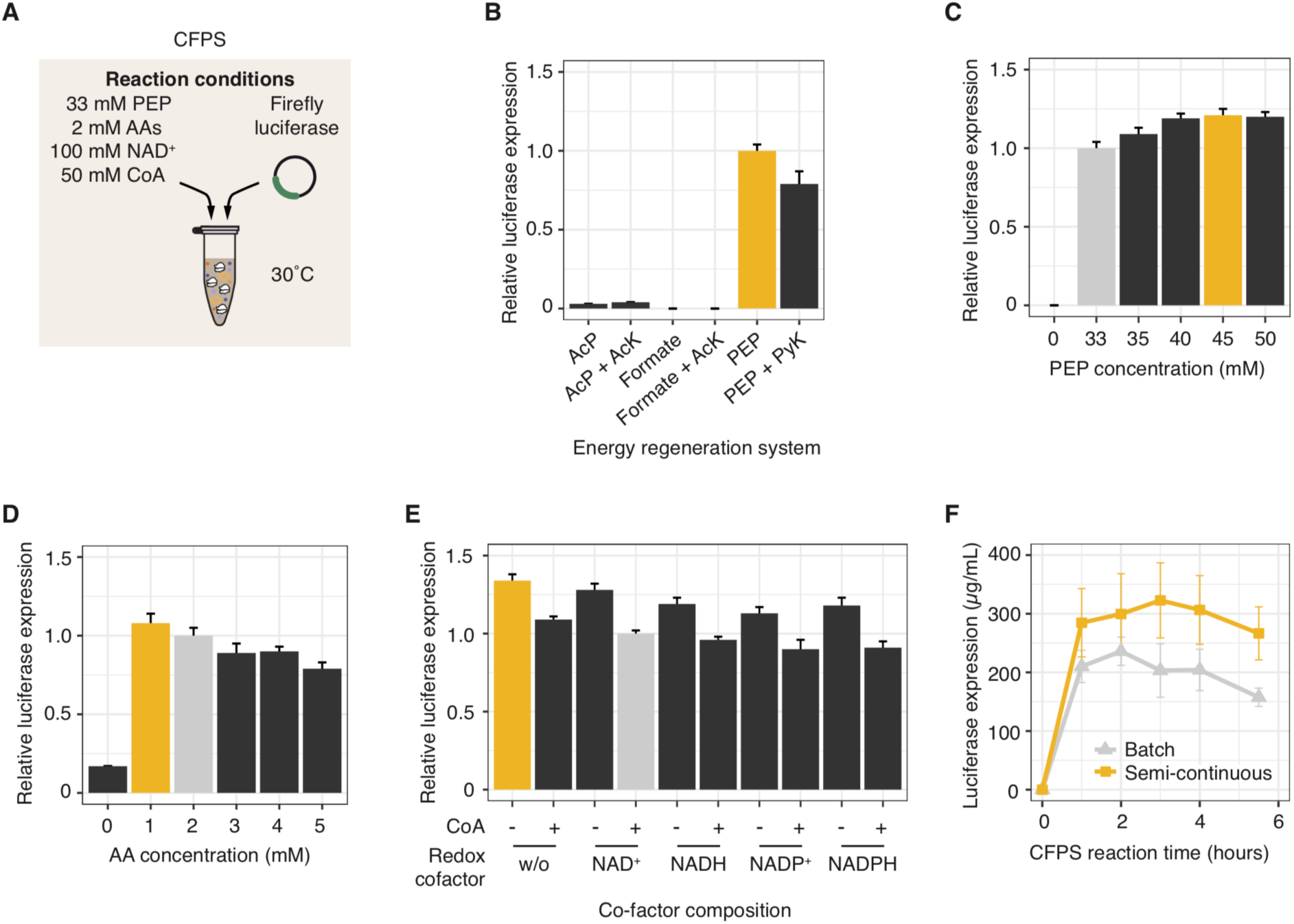
Optimization of CFE reaction conditions for *C. auto* extracts. (A) Schematic diagram of a CFE reaction depicting the concentrations of key CFE components which were step-wise adjusted for *C. auto* extracts-based CFE. **(B-H)** Relative maximum luciferase luminescence *in vitro* from *C. auto* extracts during CFE at different reagent concentrations: (B) energy regeneration system, (C) PEP concentration, (D) amino acid concentration, (E) nicotinamide dinucleotide and coenzyme A cofactor composition, (F) plasmid DNA template concentration. Light grey and yellow bars indicate previously and newly optimized condition, respectively. Maximal luciferase expressions were determined by bioluminescence and plotted as relative values compared to the previously used condition (B-E) or converted to protein yields using a luciferase standard curve (F). AA: amino acid, AcP: acetyl-phosphate, AcK: acetyl-phosphate kinase, PEP: phosphoenol pyruvate, PyK: pyruvate kinase. Data are presented as mean ± s.d. of at least three independent reactions.

First, we investigated CFE reaction temperature. To test this effect, we carried out CFE at 16 °C, 23 °C, 30 °C, and 37 °C (**Suppl. Figure S1**). We found that 16 °C and 23 °C decreased luciferase expression to 48 ± 2% and 71 ± 4%, respectively, relative to 30 °C. While CFE of luciferase at 30 °C and 37 °C increased similarly during the first 30 min, luciferase luminescence gradually decreased at 37 °C to 9 ± 1% the amount produced at 30 °C at 5.75 h. Hence, we concluded that 30 °C is the temperature optimum for *C. auto*-based CFE of luciferase, recognizing that temperature optimum may vary depending on the protein expressed.

Next, we explored secondary energy regeneration systems in *C. auto* extracts. Protein synthesis is the most energy-dependent process of exponentially growing bacterial cells, requiring ATP to be regenerated during transcription and translation. The primary source of ATP in the state-of-the-art *E. coli-*based PANOx-SP energy regeneration system (Jewett and Swartz, 2004) is phosphoenolpyruvate (PEP) conversion to pyruvate by pyruvate kinase (PyK). While this reaction occurs in *C. auto*, the Wood-Ljungdahl pathway along with acetyl-phosphate kinase (AcK) reaction is more active in generating ATP for protein synthesis (Brown et al., 2014; Kracke et al., 2016; Liew et al., 2017). Due to the difference in metabolism of *E. coli* and *C. auto*, we tested energy regeneration systems based on PEP, acetyl-phosphate (AcP), and formate, a key Wood-Ljungdahl pathway metabolite. In order to mitigate potential down-regulation or oxidative damage of the substrate’s-metabolizing enzymes in the extract due to aerobic extract preparation, we also tested supplementing 0.67 mg/mL of purified recombinant PyK with PEP and AcK with AcP and with formate. We found that almost no luciferase was expressed in the presence of substrates other than PEP (**Figure 4B**). We further investigated a range of PEP concentrations (from 0-50mM) in the “PEP+PyK” energy regeneration system (**Suppl. Figure S2A**). Interestingly, compared to PEP alone, PEP plus pyruvate kinase (PyK) reduced CFE productivity by about 20%. This inhibitory effect might be caused by the PyK storage buffer or as a result of by-products arising from the accelerated conversion of PEP to pyruvate. In addition, we see that 45 mM PEP maximized protein synthesis yields both with and without added PyK (**Suppl. Figure S2A; Figure 4C**). The maximum luciferase yield was achieved for 45 mM PEP without PyK. This was selected as the secondary energy regeneration system moving forward.

Following the optimization of the extract secondary energy source, we evaluated amino acid (AA) and co-factor concentrations. In *E. coli* extracts, supplementation of 2 mM AAs ensures adequate availability for protein synthesis and background metabolism (Martin et al.). To optimize the AA concentration for *C. auto-*based CFE, we assessed protein expression in the presence of different supplemented AA concentrations (from 0-5 mM) (**Figure 4D**). Concentrations higher than 2 mM gradually decreased the CFE yields, while reducing AAs to 1mM slightly increased luciferase expression. In addition, we found that a second supplementation of AAs after one hour of CFE had no significant effect on CFE yields (**Suppl. Figure S3**). Furthermore, NAD^+^ and CoA both have important roles in redox balancing and metabolism and are added to CFE reactions to ensure that the extract’s metabolic activity drives ATP production for protein synthesis. In contrast to *E. coli*, *C. auto* uses NADP(H) for many catabolic reactions and pyruvate oxidation to acetyl-CoA is independent of NAD(H) but relies on oxygen labile ferredoxin (Meinecke et al., 1989; Mock et al., 2015). In addition to these differences, aerobic *C. auto*-based CFE may affect the redox state and the ratio of co-factors may shift. We therefore sought to examine the impact of co-factor composition on *C. auto* extract-based CFE. We determined luciferase expression in CFE in the presence or absence of added NADP(H) or NAD(H) and with or without added CoA (**Figure 4E**). Interestingly, we found that excluding both CoA and NAD(P)(H) from the reagent mix improved luciferase expression by a third. These results together informed our selection of 1 mM AA and our decision to remove the supplementation of co-factors, which also reduces cost, going forward.

Having established concentrations for the CFE reaction buffer, we next tested the other two components of CFE: the extract and the DNA template. Increasing the extract amount has been observed to benefit other extract-based CFE systems (Li et al., 2018). We therefore tested varying volume amounts from 13-47% by volume of *C. auto* extracts on CFE. However, we did not observe any improvement in protein synthesis beyond our base case of 33% volume of extract per volume of reaction (**Suppl. Figure S4A**). We next tested whether increasing the plasmid DNA concentration from our initial 6 nM plasmid DNA would improve CFE in *C. auto* extracts as was helpful in other CFE systems (Li et al., 2017). We tested 0-30 nM of plasmid DNA and found that concentrations ≥ 15 nM increased luciferase expression by about 10-15% (**Suppl. Figure S5B**). We then tested whether linear DNA templates can be used in *C. auto* extract-based CFE. Using linear templates made by PCR avoids laborious cloning steps and can speed-up preparation time, but the template can be susceptible to exonucleases in cellular extracts, which we know have high activity in live clostridia (Nakotte et al., 1998). To test their suitability in *C. auto* extract-based CFE, we amplified the luciferase gene including its regulatory elements and additional ∼ 250 bp on the 5’- and 3’-ends from the plasmid template via PCR using standard oligonucleotide primers and with oligonucleotide primers containing phosphorothioate (PS) bonds (**Suppl. Table S1**) for increased linear template stability. Comparing CFE from reactions containing equal molarities of DNA template, we found that linear PCR products are indeed suitable templates in *C. auto* extract-based CFE (**Suppl. Figure S5A**). Using PCR products made by standard primers decreased CFE yields by only about 10%. Surprisingly, however, linear templates containing PS bonds at the 5’ and 3’ end reduced CFE yields to 50%. We also determined the optimal concentration of PCR products made by standard primers, and found it to be 33.3 nM, yielding luciferase expression comparable with the ones gained by using plasmid templates (**Suppl. Figure S5C**). This result is important because it enables a high-throughput platform where one can go from DNA sequence to protein in under 2 h.

We then explored the impact of surface area to volume ratio on the CFE reaction by changing the reaction vessel. Decreasing this ratio decreases oxygen availability and lowers the effective oxygen concentration in the reaction and thereby its availability for metabolism, which is harmful for *E. coli* extract-based CFE (Voloshin and Swartz, 2005). We tested this effect on *C. auto-*based CFE by performing 15-90 µL reactions in 1.5 mL reaction tubes and compared their luciferase expression to 40 µL reactions used previously. Increasing oxygen availability by running 15 µL reactions resulted in a ∼20% reduction in luciferase expression. However, decreasing the oxygen availability did not show significant differences (**Suppl. Figure S4B**).

Finally, we tested whether running reactions in a semi-continuous fashion, which offers substrate replenishment and byproduct (*e.g.*, inorganic phosphate) removal could further increase expression yields in *C. auto*-based CFE. To test this question, we performed CFE reactions in two compartments (complete reaction in one; reaction buffer without extract in the other) separated by a semi-permeable membrane (3.5 kDa cutoff). Small molecules can freely diffuse between both compartments, while metabolic enzymes and the translation machinery remain in the reaction compartment. We observed 37 ± 14% more active luciferase in semi-continuous reactions than in batch reactions (**Figure 4F**). Using all optimized conditions (CFE reagent concentration, extract volume, DNA template concentration) (**Suppl. Table S2**), we made 236 ± 24 µg/ ml of luciferase in batch mode and 323 ± 64 µg/ ml in semi-continuous reaction mode (**Suppl. Figure S6**).

### C. auto extract-based CFE facilitates prototyping

Upon demonstration of robust and high-yielding protein expression from our *C. auto* combined transcription and translation system, we then set out to demonstrate the potential for synthetic biology applications. In the first example, we explored the possibility for rapidly assessing and validating genetic part performance by preparing DNA templates without time-consuming cloning work. As a proof of concept demonstration for such screening capabilities, we wanted to explore (i) codon adaptation effects, (ii) the use of endogenous RNA polymerases, and (ii) expression of biosynthetic enzymes (**Figure 5A**). This is important because it addresses a key need for the development of a cell-free approach for engineering efforts in clostridia (Joseph et al., 2018).

**Figure 5:**
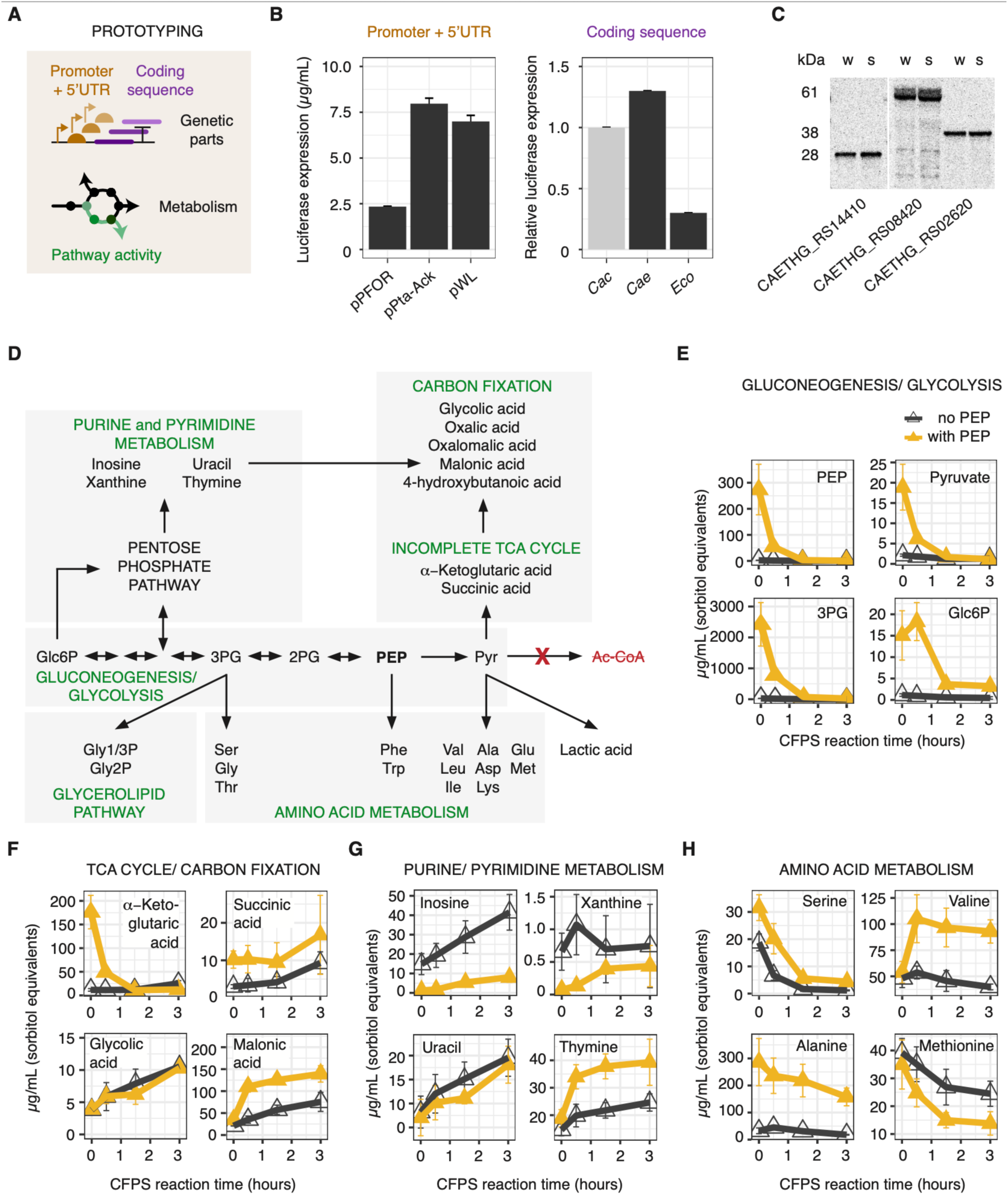
*C. auto* CFE facilitates prototyping applications towards clostridia metabolic engineering. (A) Schematic illustration of tested prototyping applications using *C. auto* CFE. (B) left panel: luciferase expression from plasmid DNA templates containing native *C. auto* promoters and a *C. auto* adapted coding sequence. Right panel: luciferase expression from PCR product templates containing coding sequences adapted for two different *Clostridium* species, *C. acetobutylicum* (Cac) and *C. auto* (Cae), and an aerobic bacterium, *E. coli* (Eco). CFE was performed using the optimized conditions. Maximal luciferase expressions were determined by bioluminescence and either plotted as luciferase yields determined by using a luciferase standard curve (left panel), or as relative values compared to the *C. auto* adapted coding sequence (right panel). Data are presented as mean ± s.d. of at least three independent reactions. (C) Autoradiography of full-length expression of recombinant native metabolic enzymes in *C. auto* CFE. CFE reactions were performed using the optimized conditions and radioactive ^14^C-Leucine (10 μM) supplemented in addition to all 20 standard amino acids. Following CFE, 4 µl CFE reaction were used for SDS-PAGE. The gels were dried and exposed for 14 days on a Storage Phosphor Screen. This image was digitally compared to the stained image that included a protein standard ladder to determine the length of synthesized proteins. w and s: whole and soluble fraction, respectively. (D) Schematic illustration of the generalized carbon flux in *C. auto* extracts during CFE reactions. (E, F, G, H) Metabolite concentration changes during CFE with or without 45 mM PEP as indicated in yellow or grey, respectively determined by GC-MS. Data are presented as mean ± s.d. of four independent reactions.

First, we compared luciferase expression using luciferase gene sequences codon-adapted for two different *Clostridium* species, *C. acetobutylicum* (*Cac*), *C. auto* (*Cae*), and *E. coli* (*Eco*), which has a significantly different global GC content (*C. auto* has a GC content of 31.1% and *E. coli* has a GC content of 50.8%) (Brown et al., 2014). We found that compared to luciferase expression from the *C. auto*-adapted sequence 20% less luciferase was expressed from a *C. acetobutylicum*-adapted one and ∼75% less from the sequence adapted for *E. coli* (**Figure 5B, right panel**). These results correlate with the predicted translation rate determined by the Salis RBS calculator (Salis et al., 2009) and with the GC content of the gene sequences (**Suppl. Figure S7**). Taken together, our data provide a proof-of-principle that the *C. auto*-based CFE systems could be used for genetic part prototyping. Second, we investigated the activity of endogenous RNA polymerases by swapping the T7 promoter and the 5’ UTR of our expression vector with three different clostridia native promoter regions (pPta-Ack, pPFOR, and pWL) that have been used for gene expression in the past (Liew et al., 2016), and compared their CFE yields. We detected luciferase expression in the range of 2-7.5 µg/mL from endogenous promoters (**Figure 5B, left panel**). As expected, the native promoter-based expression is ∼5% of the T7 promoter-based expression in *C. auto* extracts. Third, we wanted to test full-length synthesis of recombinant proteins other than luciferase. Thus, we expressed three recombinant enzymes with different protein lengths in *C. auto-*based CFE. All three enzymes were expressed in full-length, determined using an autoradiogram following incorporation of radioactive ^14^C-leucine (**Figure 6C**). In sum, our results here join an emerging wave of reports that highlight how CFE screening platforms can be used to assess if genetic designs function as expected and can express proteins in full-length.

Crude lysates are becoming an increasingly popular alternative to build biosynthetic pathways and assess their performance because they inherently provide the context of native-like metabolic networks from cytoplasmic enzymes in the lysate that remain active (Dudley et al., 2019; Karim et al., 2019b; Karim and Jewett, 2016; Kay and Jewett, 2020). We wondered the extent to which clostridia metabolism was functioning. Therefore, we set out to quantify active metabolic pathways in *C. auto* extracts. To do so, we determined the metabolome over the course of 3-hour CFE reactions with and without PEP and with and without DNA template for protein synthesis via GC-MS. We identified 44 metabolites (**Supplementary dataset**). The addition of DNA template for CFE caused only minor effects on the metabolite profiles, which has been seen previously in *E. coli* cell-free systems (Karim et al., 2018), leading us to pool together the sample sets identical in PEP treatment and CFE reaction time. We split the detected metabolites into specific anabolic and catabolic reactions based on generalized carbon flux in *C. auto* extracts during CFE (**Figure 5D**). For most identified metabolites, we detected concentration changes during the cell-free reaction, indicating metabolic activity of their corresponding biosynthesis and degradation pathways (**Figure 5E-H**). We observed large-scale effects when comparing metabolites from reactions with and without. For instance, PEP addition immediately increased the concentrations of glycolysis/ gluconeogenesis intermediates 3-phosphoglyceric acid (3PG), 2-phosphoglyceric acid, and glucose 6-phosphate (Glc6P) (**Figure 5E**). Additionally, several organic acids were up-regulated, including metabolites involved in tricarboxylic acid (TCA) cycle and carbon fixation into biomass, such as α-keto-glutaric acid, succinic acid, glycolic acid and malonic acid (**Figure 5F**). Metabolites that were depleted in CFE reactions containing PEP included the purine and pyrimidine pathway intermediates inosine, xanthine and uracil (**Figure 5G**) and the amino acid methionine (**Figure 5H**). In summary, we observed metabolites of glycolysis/gluconeogenesis and associated pathways, including nucleotide synthesis, incomplete TCA cycle, carbon fixation, amino acid and glycerolipid pathway. We did not observe carbon flux towards acetyl-coA and associated pathways. This observation indicates that the enzyme converting pyruvate to acetyl-CoA, pyruvate:ferredoxin oxidoreductase (PFOR), is inactive in aerobic *C. auto* extracts as described for other clostridia (Meinecke et al., 1989). Together, our results suggest that the developed *C. auto* cell-free system could indeed be used to test libraries of genetic parts and study metabolic pathways.

## DISCUSSION

In this work, we describe the development of a robust, high-yielding CFE system derived from the non-model and anaerobic bacterium *C. auto.* To do so, we optimized the extract lysis preparation procedure, streamlined the extract processing steps, and optimized cell-free reaction conditions. We found *C. auto*-derived CFE requires different conditions than *E. coli*-derived CFE. Surprisingly, *C. auto* CFE requires unusually high magnesium concentrations. Our final system was able to produce more than 230 µg/mL of luciferase within a 3-hour batch reaction, making it one of the more productive cell-free systems developed to date. Indeed, this batch CFE yield is higher than that of most other systems derived from model and non-model organisms such as rabbit reticulocytes (Anastasina et al., 2014), archaea (Endoh et al., 2008, 2007, 2006), yeast (Gan and Jewett, 2014; Hodgman and Jewett, 2013), insects (Ezure et al., 2010), *Bacillus subtilis* (Kelwick et al., 2016)*, Bacillus megaterium* (Moore et al., 2018) and *Streptomyces* (Li et al., 2018, 2017). Performing semi-continuous reactions, we increased yields to more than 320 µg/mL within 4 hours. Only CFE systems derived from CHO cell (Martin et al., 2017), *V. natriegens* (Des Soye et al., 2018), wheat germ (Harbers, 2014) and *E. coli* (Caschera and Noireaux, 2014; Des Soye et al., 2019) have been demonstrated to be more productive. We anticipate that our optimization workflow can pave the way for development of CFE systems for clostridia species including solventogenic or cellulolytic clostridia but also medical relevant clostridia. Further optimizing CFE reaction conditions could help prolong the CFE reaction duration and thereby further increase protein yields and development of an anaerobic system may mimic the cellular environment of clostridia to facilitate natural metabolism.

In addition to showing high yielding CFE, we demonstrated that the *C. auto*-based CFE system is compatible with PCR amplicon as expression templates with minimal purification required. We also showed our system’s capability as a gene regulatory screening platform by monitoring gene expression from a library with different codon optimized sequences. Importantly, our system also allows prototyping of native promoters that need to be recognized by the endogenous transcription machinery. This is a particularly powerful feature of our system. The most commonly used promoters for clostridia metabolic engineering originate from a few strains and are often not transferrable to non-native hosts. Being able to characterize promoter parts and to test adjustments rapidly and in high-throughput may have a significant impact on clostridia metabolic engineering.

Looking forward, we believe the CFE platform will serve as a useful “toolbox” for clostridia metabolic engineering and help accelerate strain engineering efforts. A key enabling feature of this toolbox is that cellular metabolism is active from native pathways in the lysate. By integrating automatic liquid handling systems, data-driven AI approaches should expedite screening and prototyping efforts in clostridia for a variety of synthetic biology applications.

## Materials & Methods

### Strains and plasmid constructs

*Clostridium autoethanogenum* DSM 19630, a derivate of type strain DSM10061 was used in this study (Heijstra et al., 2016). The gene sequences and oligonucleotides used in this study are listed in the **Supplemental Materials and Table S1, respectively**.

Codon-adapted luciferase genes for CFE were synthesized by IDT, cloned into the pJL1 plasmid using Gibson assembly and confirmed by Sanger sequencing by ACGT, Inc. Kanamycin (50 μg/ mL) was used to maintain pJL1-based plasmids. *C. auto* endogenous promoters of phosphotransacetylase-actetate kinase operon (pPta-Ack; CAETHG_RS16490), pyruvate:formate oxidoreductase (pPFOR; CAETHG_RS14890) and Wood-Ljungdahl cluster (pWL; CAETHG_RS07860) were amplified from a plasmid where the respective sequences have been amplified from the genome and cloned into a pMTL82250 vector reporter plasmid (Nagaraju et al., 2016) and cloned in place of the T7 promoter region in the pJL1-LucCae construct using Gibson assembly and confirmed by Sanger sequencing by ACGT, Inc.

### Cell culture and harvest

Fermentations with *C. auto* were carried out in 10-L bioreactors with a working volume of 6 L at 37 °C and CO-containing gas (50% CO, 10% H_2_, 20% CO_2_, 20% N_2_) as sole energy and carbon source at a bacterial growth rate near 1 day^-1^ as described earlier (Wang et al., 2013). Prior to harvest of the cells, the pH of the culture was adjusted to pH 6 with K_2_CO_3_. Five liters of culture were collected on ice. The culture was divided between 1-L centrifuge bottles and cells pelleted at 5,000 x g for 10 min. The supernatant was decanted, and residual liquid removed. The pellets were resuspended in ∼300 mL of 50 mM K_2_PO_4_, pH 7.5. Resuspensions were transferred to 50-mL-Falcon-tubes and cells pelleted at 5,000 x g for 15 min. Supernatants were discarded and the pellets immediately frozen on liquid N_2_ and stored at −80 °C.

### Extract preparation

Cell pellets were thawed and suspended in 0.33 mL of S30 buffer (10 mM Tris(CH_3_COO) (pH 8.2), 14 mM Mg(CH_3_COO)_2_, 10 mM K(CH_3_COO), 4 mM DTT) per gram of wet cell mass. The cell suspension was transferred as 1 mL aliquots into 1.5 mL microtubes. Using a Q125 Sonicator (Qsonica, Newtown, CT, USA) with 3.175 mm diameter probe at a 20 kHz frequency and 50% amplitude, cells were lysed for several cycles of 10s ON/10s OFF until final input energy was reached. Samples were kept in an ice-water bath during sonication to minimize potential heat denaturation arising from sonication. For each 1 mL cell suspension aliquot, the input energy was ∼70 Joules/ sonication cycle. Subsequently, lysates were centrifuged at 12,000 × *g* at 4 °C for 10 min, supernatants collected, flash-frozen in liquid nitrogen, and stored at −80 °C until use. For run-off reactions, the supernatant of the first clarifying spin was transferred to a new tube, incubated at 37 °C for 45 min or 80 min, cleared by centrifugation at 12,000 x *g* at 4 °C for 10 min, supernatants collected, flash-frozen in liquid nitrogen, and stored at −80 °C until use. Dialysis was performed using Slide-A-Lyzer™ Dialysis Cassettes with a 3.5 kDa cut-off (Thermo Scientific, Rockford, IL, USA). Extracts were dialyzed three times with 150 mL S30 buffer per mL extract for 45 min at 4 °C, and subsequently cleared by centrifugation at 12,000 × *g* at 4 °C for 10 min. Supernatants were collected, flash frozen in liquid nitrogen, and stored at −80 °C until use.

### CFE reaction

A modified PANOx-SP system was utilized for CFE reactions. Briefly, if not stated otherwise, in a 1.5 mL microtube 40-60 μL CFE reactions were prepared by mixing the following components: 1.2 mM ATP; 0.85 mM each of GTP, UTP, and CTP; 34 μg/ mL folinic acid; 170 μg/mL of *E. coli* tRNA mixture; 16 μg/mL T7 RNA polymerase; 2 mM for each of the 20 standard amino acids; 0.33 mM nicotinamide adenine dinucleotide (NAD); 0.27 mM coenzyme-A (CoA); 1.5 mM spermidine; 1 mM putrescine; 4 mM sodium oxalate; 8 mM magnesium glutamate; 10 mM ammonium glutamate; 130 mM potassium glutamate; 57 mM HEPES (pH 7.2); 33 mM phosphoenolpyruvate (PEP), and 33% (v/v) of cell extract. Unless noted otherwise, synthesis of specific products was initiated by adding 6 nM of pJL1 template plasmid encoding the gene of interest to each reaction, and each CFE reaction was incubated at 30 °C. Because individual reagent concentrations were optimized throughout the study, their determined optimal values were used for all reactions from that point onward. *E. coli* total tRNA mixture (from strain MRE600) and PEP was purchased from Roche Applied Science (Indianapolis, IN, USA); ATP, GTP, CTP, UTP, 20 amino acids and other materials were purchased from Sigma (St. Louis, MO, USA) without further purification. T7RNAP was purified in house as described previously (Swartz et al., 2004).

### Quantification of active luciferase

Luciferase expression in CFE was determined using the ONE-Glo Luciferase Assay System (Promega, Madison, WI, USA), a Synergy 2 plate reader (BioTek, Winooski, VT, USA), and 96-well half area white plates (Costar 3694; Corning, Corning, NY). The assay was performed using 4 µl CFE reaction mixed with 30 µl of luciferase assay buffer. Luminescence was detected every 3 min over a 30 min period using a BioTek Synergy 2 plate reader (Winooski, VT, USA). The maximum amount of relative light units (RLUs) was recorded for each reaction. RLUs were then converted into µg/ mL amounts using a linear standard curve determined using radioactively labelled luciferase. For this, CFE reactions were performed with radioactive ^14^C-Leucine (10 μM) supplemented in addition to all 20 standard amino acids. Radioactively labelled protein samples were then precipitated using trichloroacetic acid (TCA) and their radioactive counts measured by liquid scintillation using a MicroBeta2 (PerkinElmer, Waltham, MA) to quantify soluble and total luciferase yields as previously reported (Jewett et al., 2008; Jewett and Swartz, 2004).

### Semi-continuous CFE reaction

90 μL CFE semi-continuous reactions were performed using 3.5 kDa MWCO 96-well plate dialysis cassettes (Thermo Scientific, Rockford, IL, USA) in 2 mL microcentrifuge tubes with 1.4 mL dialysis buffer solution. Reactions were incubated in an Eppendorf Thermomixer C at 30 °C and 600 rpm and compared to a 60 μL batch reaction performed under the same conditions.

### Gas chromatography-mass spectrometry (GC-MS)

Clostridia CFE reaction samples were analyzed by GC-MS. In brief, samples stored at −80 °C prior to analysis were thawed and centrifuged at 12,000 rpm at 4 °C for 15 minutes. An aliquot of 5 µl was transferred to a vial containing 10 µl of sorbitol (1 mg/ml aqueous solution) used as internal standard and then dried under a stream of N_2_. Dried samples were dissolved in 250 µl of silylation-grade acetonitrile followed by addition of 250 µl of N-methyl-N- (trimethylsilyl)trifluoroacetamide (MSTFA) with 1% trimethylchlorosilane (TMCS) (Thermo Scientific, Bellefonte, PA) and heated for 1 hr at 70 °C to generate trimethylsilyl derivatives. After 2 days, 1 µl aliquots were injected into an Agilent Technologies 7890A GC coupled to a 5975C inert XL MS fitted with an RTX-5MS (5% diphenyl/ 95% dimethyl polysiloxane) 30 m × 250 μm × 0.25 μm film thickness capillary column with a 5 m Integra-Guard column. Gas flow was 1.0 ml per minute and the injection port was configured for spitless injection. The initial oven temperature was 50 °C with a 2-minute hold followed by a temperature ramp of 20 °C per minute to 325 °C and held for another 11.5 minutes. The MS was operated in standard electron impact (70 eV) ionization mode. The injection port, MS transfer line, MS source, and MS quad temperatures were 250 °C, 300 °C, 230 °C, and 150 °C respectively. A large user-created database and the commercially available Wiley Registry 10th Edition combined with the NIST 14 mass spectral database were used to identify metabolites of interest. Peaks were quantified by using extracted-ion chromatograms (EIC) rather than total ion current chromatograms, utilizing a key selected ion characteristic m/z fragment, to minimize co-eluting metabolites. The EIC was scaled back to TIC using predetermined scaling factors and quantification was based on area integration and normalized to the quantity of internal standard recovered, the volume of sample processed, the derivatization volume and injection volume.

### Autoradiography

Autoradiography was used to determine the quality of synthesized metabolic enzymes synthesized in *C. auto* CFE. CFE reactions were performed with radioactive ^14^C-Leucine (10 μM) supplemented in addition to all 20 standard amino acids. Following 3.5 hrs incubation, 4 µl CFE reaction was loaded onto a NuPAGE 4–12% Bis–Tris Gel (Life Technologies, Carlsbad, CA, USA) following the manufacturer’s instructions. The NuPAGE gels were stained with InstantBlue (Expedeon, Cambridgeshire, UK). The gels were dried and exposed for 14 days on a Storage Phosphor Screen (GE Healthcare Biosciences, Chicago, IL, USA) and imaged with a Typhoon FLA 7000 (GE Healthcare Biosciences). This image was digitally compared to the stained image that included a protein standard ladder to determine the length of synthesized proteins.

## Author Contribution

A.K. and M.C.J. designed the experiments. A.P.M. and M.K. generated *C. auto* cells. A.K., G.A.R., N.L.E., Z.K.Y., T.J.T. conducted and analyzed experiments. A.K. and M.C.J. wrote the manuscript. M.C.J. supervised the research.

## Acknowledgements

The authors thank Ashty Karim (ORCID # 0000-0002-5789-7715), Lauren Clark, and Ben De-Soye for helpful comments on the manuscript and Ava Rhule-Smith, Jim Daleiden, Monica MacDonald, and Steven Glasker for their help growing the *C. auto* cultures and Robert Nogle for help with subcloning the promoter regions. This work was supported by the U.S. Department of Energy, Office of Biological and Environmental Research in the DOE Office of Science under Award Number DE-SC0018249. M.C.J. also thanks the David and Lucille Packard Foundation and the Camille-Dreyfus Teacher-Scholar Program for their generous support. We also thank the following investors in LanzaTech’s technology: BASF, CICC Growth Capital Fund I, CITIC Capital, Indian Oil Company, K1W1, Khosla Ventures, the Malaysian Life Sciences, Capital Fund, L. P., Mitsui, the New Zealand Superannuation Fund, Novo Holdings A/S, Petronas Technology Ventures, Primetals, Qiming Venture Partners, Softbank China, and Suncor.

## Supplementary Information

### Supplementary Tables

**Table S1.**
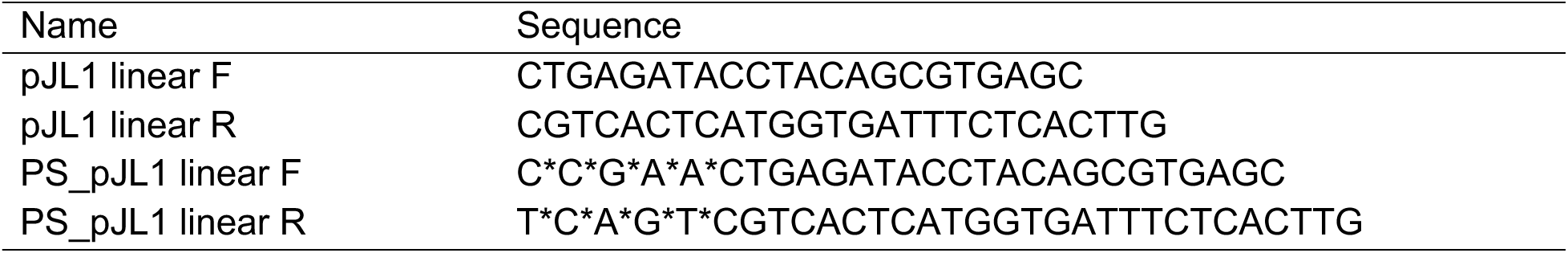
Oligonucleotide sequences.

**Table S2.** Optimized cell-free protein synthesis conditions.

| Reagent | Final concentration or volume |
| --- | --- |
| Mg(Glu)<sub>2</sub> | 24 mM |
| Amino acids (all 20) | 1 mM |
| PEP | 45 mM |
| NAD<sup>+</sup> | 0 mM |
| CoA | 0 mM |
| Plasmid DNA | 15 nM |
| C. auto extract | 5 µL |

### Supplementary Figures

**Figure S1.**
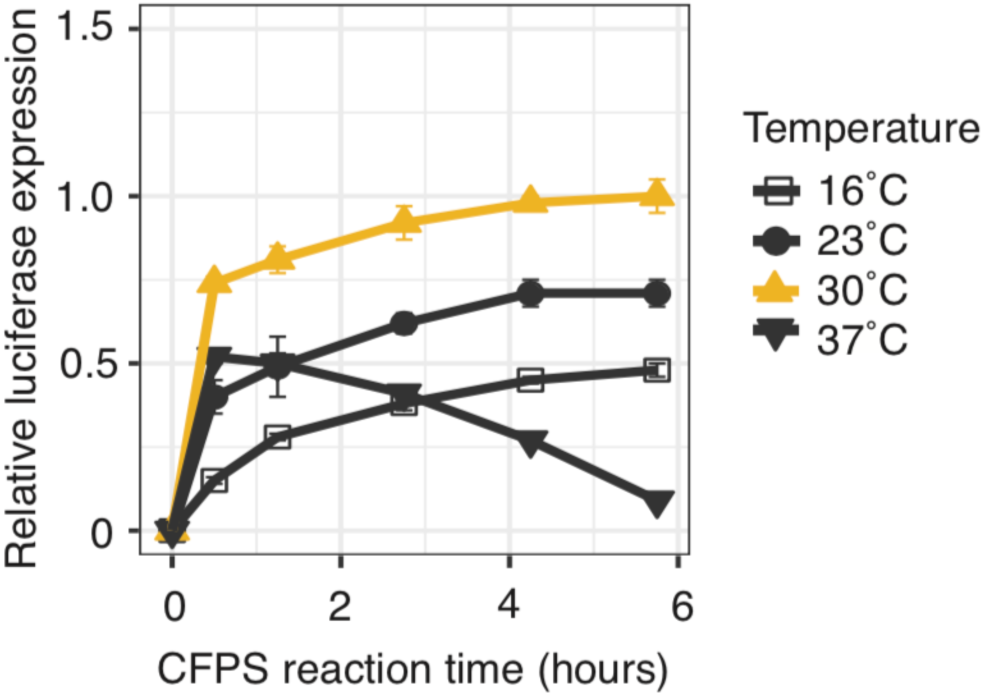
Effect of temperature on CFE with *C. auto* extracts. Relative luciferase expression in CFE at different reaction temperatures. *C. auto* cell pellets were resuspended in S30 buffer, lysed by sonication at 350 J, clarified by centrifugation at 12,000 x g, dialyzed against S30 buffer 3 times for 45 min at 4 °C, clarified again by centrifugation at 12,000 x g and used for CFE at indicated temperatures. Luciferase expression was determined by bioluminescence and plotted as relative values compared to the maximal luciferase expression at 30 °C. Yellow line highlights optimal condition. Data are presented as mean ± s.d. of three independent reactions.

**Figure S2.**
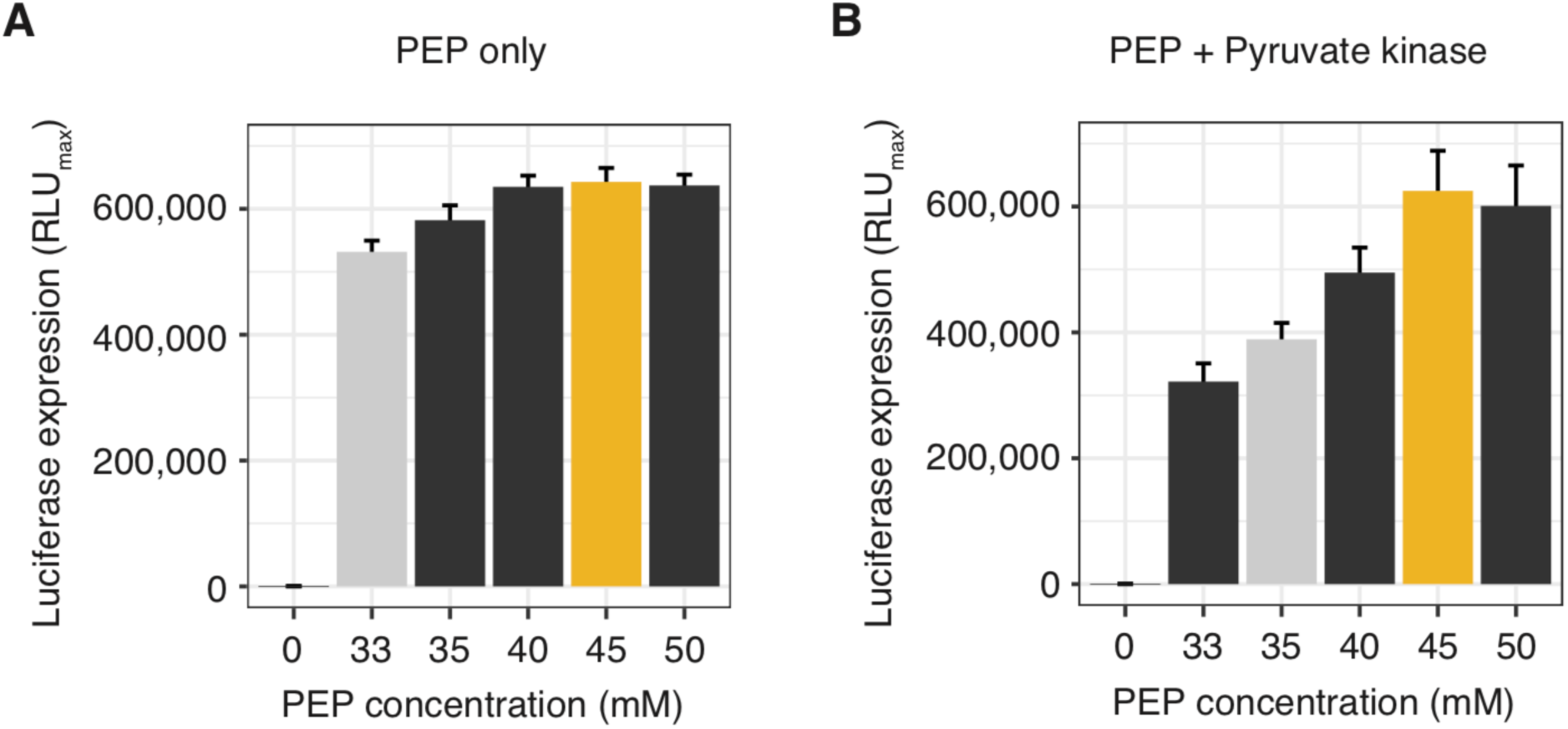
Effect of phosphoenolpyruvate (PEP) on CFE with *C. auto* extracts. Maximal luciferase expression during CFE (A) at different concentrations of PEP and (B) at different concentrations of PEP in the presence of pyruvate kinase (PyK). CFE was performed with indicated PEP concentrations at 30°C. Luciferase expression was determined by bioluminescence. Light grey and yellow bars indicate previously used and newly optimized condition, respectively. Data are presented as mean ± s.d. of three independent reactions.

**Figure S3.**
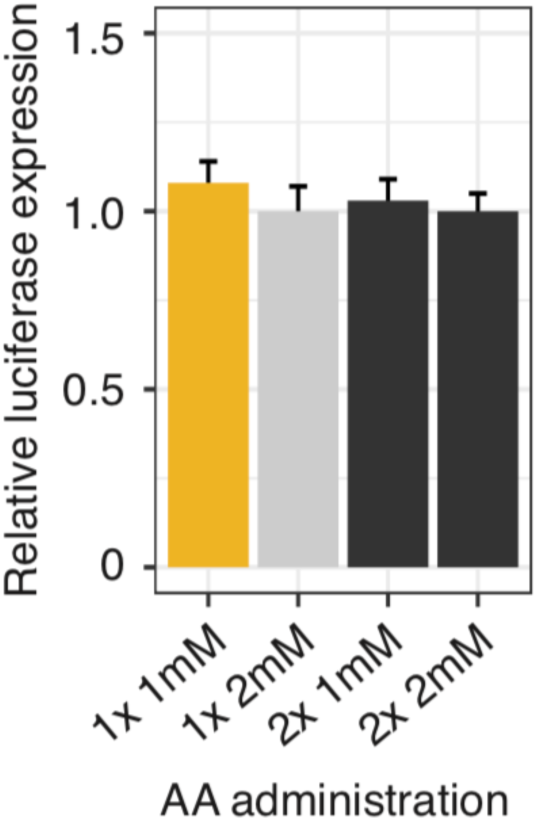
The impact of amino acid concentration (AA) on CFE with *C. auto* extracts. Relative luciferase expression under different AA administration regimes. CFE reactions were performed with 45 mM PEP and indicated AA concentrations, either 1 mM or 2 mM (1x). In addition to batch reactions with different starting AA concentrations, reactions were supplemented after 1 h of CFE (2x). Luciferase expression was determined by bioluminescence and plotted as relative values compared to the previously used condition. Light grey and yellow bars indicate previously and newly optimized condition, respectively. Data are presented as mean ± s.d. of three independent reactions.

**Figure S4.**
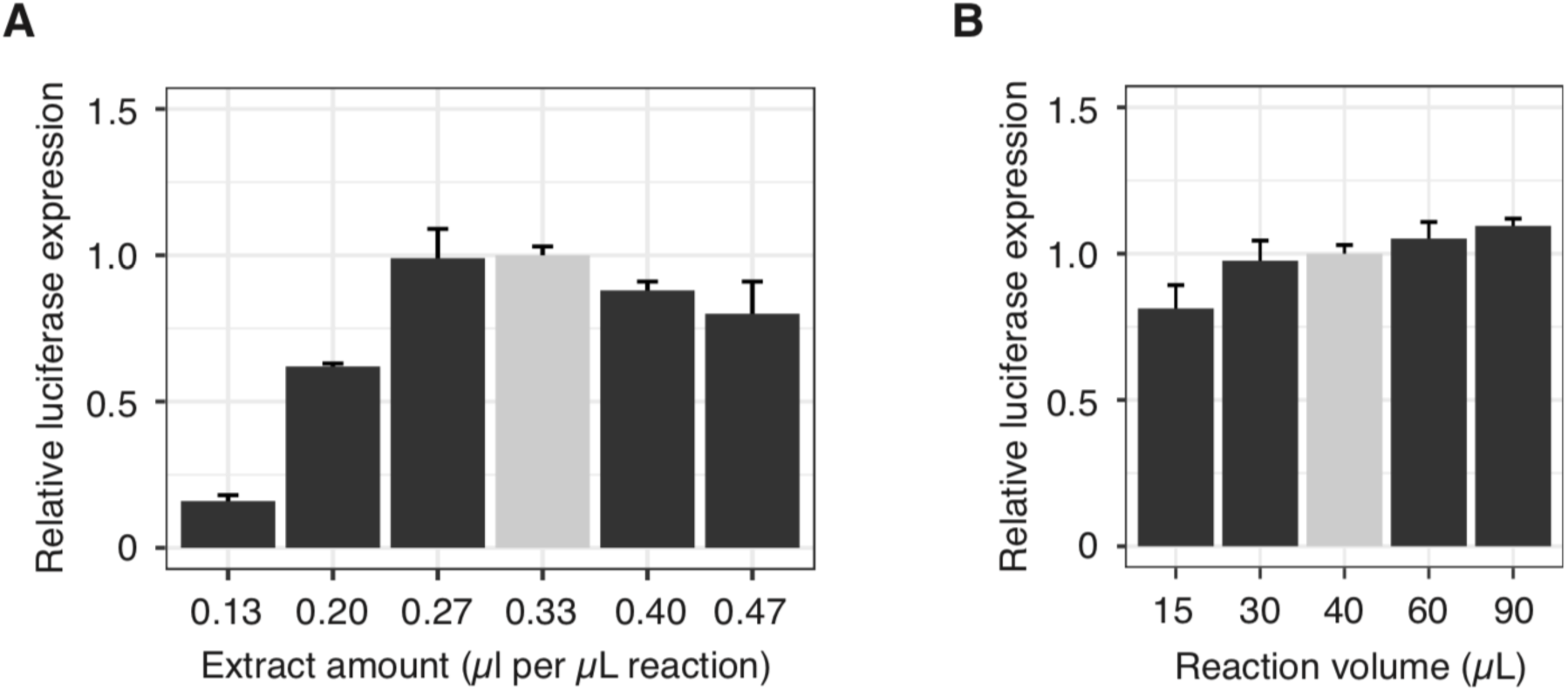
Effect of extract amount and oxygen availability on CFE with *C. auto* extracts. (A) Relative luciferase expression in CFE reactions containing different amounts of extract per µl CFE reaction. CFE reactions were set up as 40 µl batch reactions. (B) Relative luciferase expression in CFE batch reactions set up at indicated volumes in 1.5 ml reaction tubes. Luciferase expression was determined by bioluminescence and plotted as relative values compared to the luciferase expression of the previously used condition (light grey bar). Data are presented as mean ± s.d. of three independent reactions.

**Figure S5.**
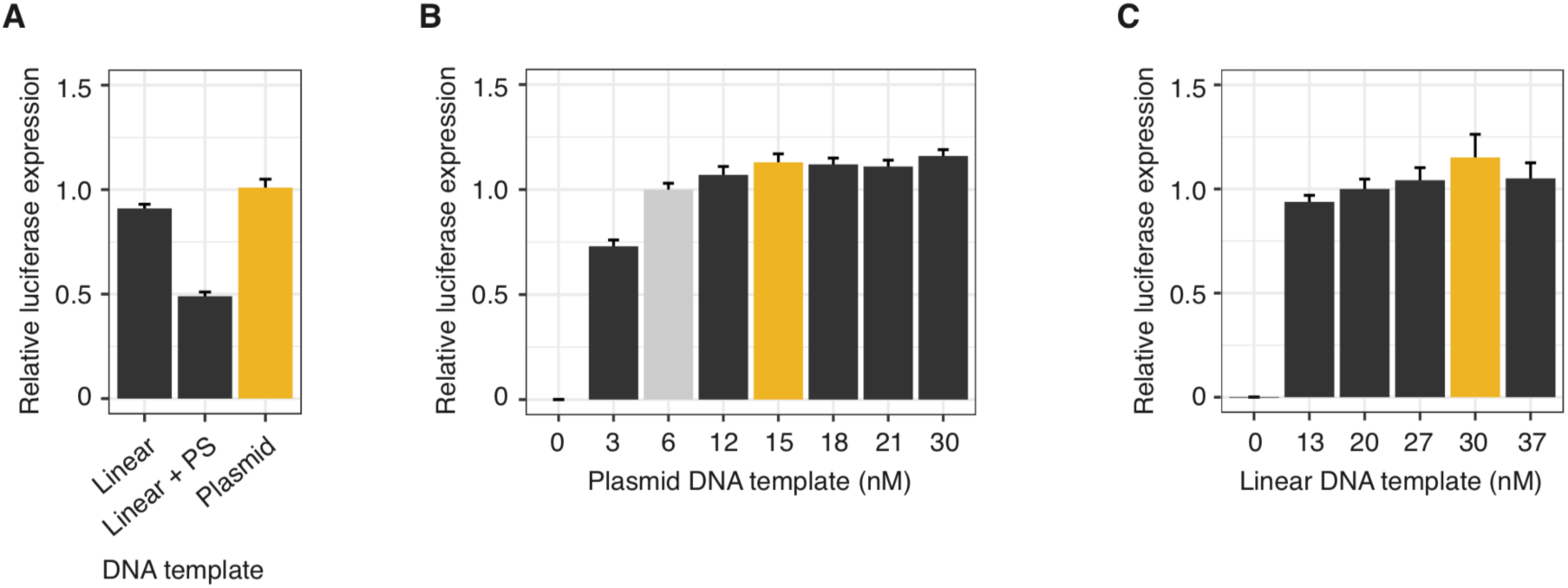
Effect of DNA template type and concentration on CFE with *C. auto* extracts. Relative luciferase expression in CFE reactions containing (A) different DNA template types, and different concentrations of (B) plasmid DNA, and (C) linear DNA. Plasmid DNA was midi-prepped and additionally cleaned-up by ethanol precipitation. Linear templates were prepared by PCR and cleaned using a PCR purification kit and additional ethanol precipitation. Luciferase expression was determined by bioluminescence and plotted as relative values compared to the luciferase expression of the previously used condition (light grey bar). Yellow bar indicates newly optimized condition. Data are presented as mean ± s.d. of three independent reactions.

**Figure S6.**
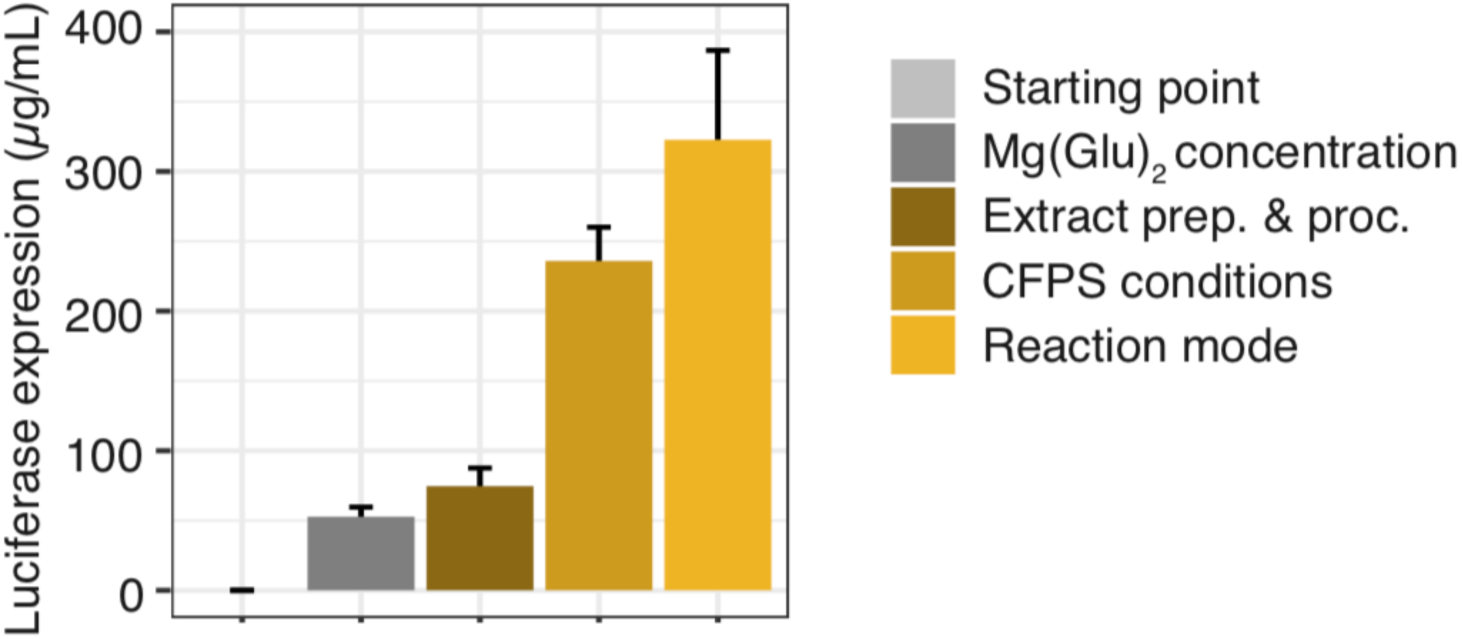
Summary of the development, optimization, and improvements of C. auto CFE. Shown are step-wise and cumulative improvements of luciferase expression yields in *C. auto* CFE. Extract prep. & proc.: extract preparation and processing. Data are presented as mean ± s.d. of at least three independent reactions.

**Figure S7.**
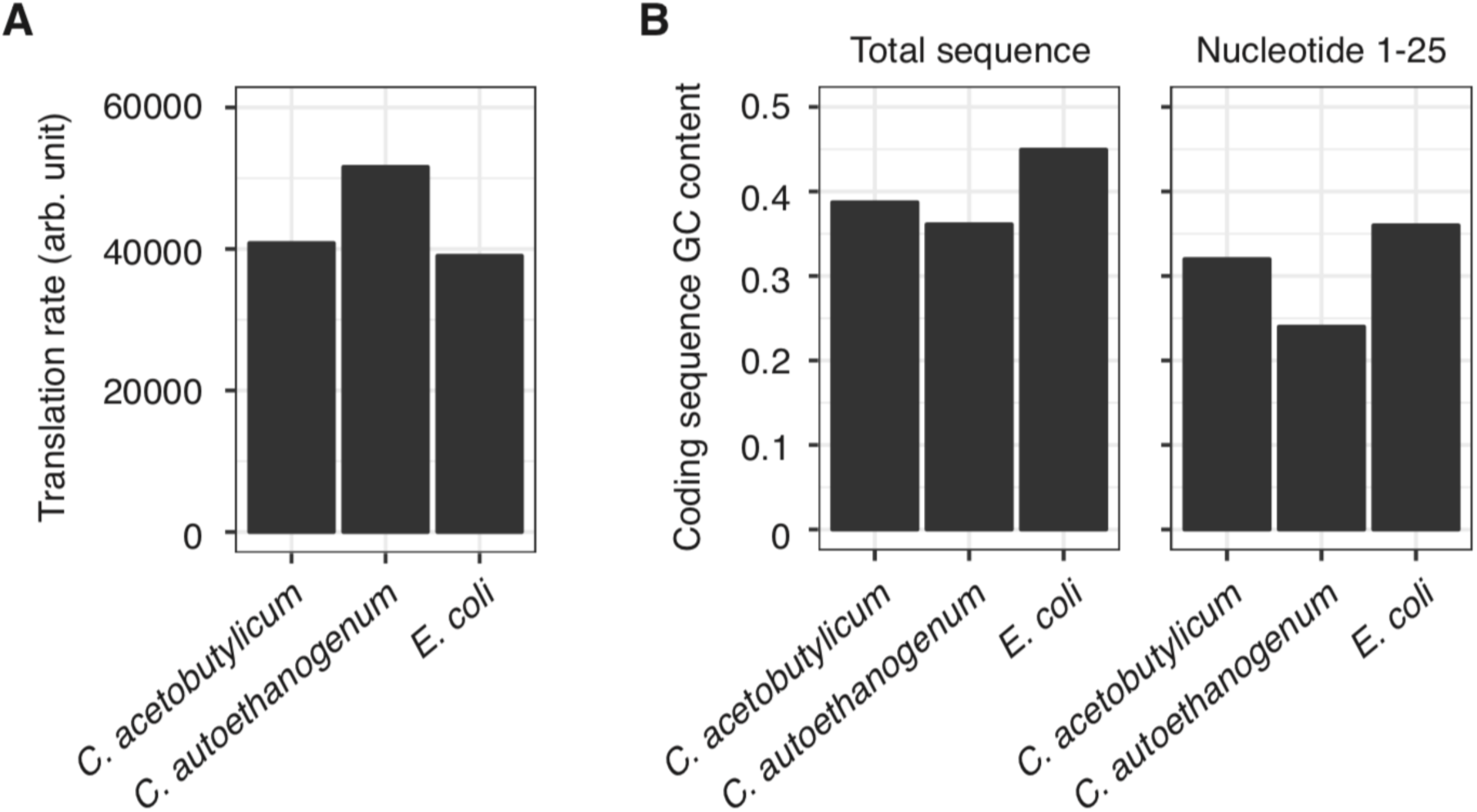
Analysis of three different luciferase coding sequences. (A) Predicted translation rate determined by an RBS calculator (Salis et al., 2009). (B) GC content of luciferase coding sequences.

## Supplemental Materials

### Sequences

#### Promoter + 5’UTRs

##### phosphotransacetylase-acetate kinase operon (pPta-Ack; CAETHG_RS16490) promoter

GGCCGCAATATGATATTTATGTCCATTGTGAAAGGGATTATATTCAACTATTATTCCAGTTACGTTCATAGAAATTTTCCTTTCTAAAATATTTTATTCCATGTCAAGAACTCTGTTTATTTCATTAAAGAACTATAAGTACAAAGTATAAGGCATTTGAAAAAATAGGCTAGTATATTGATTGATTATTTATTTTAAAATGCCTAAGTGAAATATATACATATTATAACAATAAAATAAGTATTAGTGTAGGATTTTTAAATAGAGTATCTATTTTCAGATTAAATTTTTGATTATTTGATTTACATTATATAATATTGAGTAAAGTATTGACTAGCAAAATTTTTTGATACTTTAATTTGTGAAATTTCTTATCAAAAGTTATATTTTTGAATAATTTTTATTGAAAAATACAACTAAAAAGGATTATAGTATAAGTGTGTGTAATTTTGTGTTAAATTTAAAGGGAGGAAATGAACATGAAACAT

##### pyruvate:formate oxidoreductase (pPFOR; CAETHG_RS14890) promoter

GCAAAATAGTTGATAATAATGCAGAGTTATAAACAAAGGTGAAAAGCATTACTTGTATTCTTTTTTATATATTATTATAAATTAAAATGAAGCTGTATTAGAAAAAATACACACCTGTAATATAAAATTTTAAATTAATTTTTAATTTTTTCAAAATGTATTTTACATGTTTAGAATTTTGATGTATATTAAAATAGTAGAATACATAAGATACTTAATTTAATTAAAGATAGTTAAGTACTTTTCAATGTGCTTTTTTAGATGTTTAATACAAATCTTTAATTGTAAAAGAAATGCTGTACTATTTACTGTACTAGTGACGGGATTAAACTGTATTAATTATAAATAAAAAATAAGTACAGTTGTTTAAAATTATATTTTGTATTAAATCTAATAGTACGATGTAAGTTATTTTATACTATTGCTAGTTTAATAAAAAGATTTAATTATATACTTGAAAAGGAGAGGAATCCAT

##### Wood-Ljungdahl cluster (pWL; CAETHG_RS07860) promoter

AGATAGTCATAATAGTTCCAGAATAGTTCAATTTAGAAATTAGACTAAACTTCAAAATGTTTGTTAAATATATACCAAACTAGTATAGATATTTTTTAAATACTGGACTTAAACAGTAGTAATTTGCCTAAAAAATTTTTTCAATTTTTTTTAAAAAATCCTTTTCAAGTTGTACATTGTTATGGTAATATGTAATTGAAGAAGTTATGTAGTAATATTGTAAACGTTTCTTGATTTTTTTACATCCATGTAGTGCTTAAAAAACCAAAATATGTCACATGCAATTGTATATTTCAAATAACAATATTTATTTTCTCGTTAAATTCACAAATAATTTATTAATAATATCAATAACCAAGATTATACTTAAATGGATGTTTATTTTTTAACACTTTTATAGTAAATATATTTATTTTATGTAGTAAAAAGGTTATAATTATAATTGTATTTATTACAATTAATTAAAATAAAAAATAGGGTTTTAGGTAAAATTAAGTTATTTTAAGAAGTAATTACAATAAAAATTGAAGTTATTTCTTTAAGGAGGGAATTATTCAT

#### Luciferase coding sequences

##### *C. acetobutylicum* codon-optimized

ATGGAGGATGCAAAGAATATTAAGAAAGGTCCAGCTCCTTTCTACCCTTTAGAAGACGGAACTGCTGGTGAGCAATTACACAAGGCAATGAAGAGGTATGCATTAGTACCAGGAACTATAGCTTTTACTGATGCTCATATTGAAGTAAATATAACATATGCAGAATACTTTGAGATGTCTGTGAG GCTTGCAGAAGCAATGAAAAGATATGGATTAAATACTAACCACAGGATAGTGGTTTGTTCTGAAAACAGCTTACAATTCTTCATGCCAGTTCTTGGAGCATTATTCATTGGAGTTGCTGTGGCTCCAGCAAATGACATTTACAACGAGAGGGAGTTGTTAAATTCAATGAATATTAGTCAACCTACTGTAGTGTTCGTTTCTAAGAAGGGACTTCAGAAAATTCTAAACGTGCAAAAAAAGCTACCAATTATTCAAAAGATAATAATTATGGACTCAAAAACTGATTACCAAGGATTCCAGAGCATGTATACTTTTGTTACATCTCATCTACCACCAGGTTTTAATGAGTATGATTTCGTGCCAGAAAGCTTTGACAGAGATAAGACAATAGCTTTGATTATGAACAGCTCAGGATCTACAGGACTACCTAAGGGTGTGGCTCTACCTCATAGGACTGCTTGCGTTAGGTTTAGTCATGCAAGGGACCCTATATTTGGAAATCAAATTATTCCTGATACTGCAATACTATCAGTTGTACCATTTCATCACGGATTCGGTATGTTCACAACATTGGGATATCTTATATGCGGTTTTAGAGTGGTACTTATGTATAGGTTCGAGGAAGAACTTTTTTTAAGGAGTCTACAAGACTACAAAATACAATCAGCATTGTTAGTGCCAACATTATTTAGTTTTTTCGCTAAAAGCACACTTATAGATAAATACGACTTGTCTAACCTACACGAAATAGCAAGCGGTGGAGCTCCTTTATCTAAAGAGGTAGGAGAAGCTGTTGCAAAAAGATTCCACTTACCTGGTATAAGACAGGGTTATGGATTGACAGAAACAACATCAGCAATTTTGATTACTCCAGAGGGAGATGACAAGCCAGGTGCTGTAGGAAAGGTGGTACCATTCTTTGAAGCTAAAGTAGTAGACTTGGACACAGGAAAAACTCTAGGTGTGAATCAGAGAGGAGAACTTTGCGTAAGGGGACCAATGATAATGTCAGGTTATGTAAATAATCCAGAGGCAACAAATGCACTTATAGATAAAGATGGTTGGTTGCACAGCGGAGATATAGCTTACTGGGATGAGGACGAACATTTTTTTATTGTGGACAGGCTTAAAAGTTTGATTAAATACAAAGGTTACCAGGTGGCACCAGCTGAGCTTGAATCTATATTGCTTCAACACCCAAATATTTTTGATGCTGGTGTAGCAGGTCTTCCTGATGATGATGCAGGAGAGCTTCCAGCTGCTGTAGTAGTATTAGAGCACGGAAAGACAATGACAGAAAAGGAAATAGTGGACTACGTTGCATCTCAGGTTACAACTGCAAAGAAGCTAAGGGGAGGTGTTGTTTTTGTAGACGAAGTTCCTAAAGGATTGACAGGAAAGTTGGACGCTAGGAAGATTAGAGAGATTCTAATAAAAGCTAAGAAGGGTGGTAAAAGTAAGTTATAG

##### *C. autoethanogenum* codon-optimized

ATGGAAGATGCAAAAAATATAAAGAAAGGACCAGCACCATTCTATCCACTTGAAGACGGAACAGCAGGAGAACAGCTACATAAAGCAATGAAAAGATATGCACTTGTACCGGGAACAATAGCTTTTACTGATGCTCACATAGAAGTAAACATAACCTATGCGGAATATTTTGAAATGTCAGTAAGATTGGCAGAGGCAATGAAAAGATATGGATTAAATACAAATCATAGAATAGTAGTGTGTAGTGAAAACAGCTTGCAGTTTTTTATGCCTGTACTTGGTGCTTTATTCATAGGTGTAGCAGTAGCACCAGCTAATGATATTTATAATGAACGTGAGCTTTTAAATTCTATGAATATAAGTCAGCCAACTGTAGTATTTGTTTCAAAGAAAGGTTTGCAGAAGATTTTGAATGTTCAAAAGAAATTGCCTATAATTCAAAAAATAATAATTATGGATTCTAAGACAGATTATCAGGGATTCCAGTCTATGTATACATTCGTAACATCTCATCTTCCCCCGGGATTTAATGAATATGACTTCGTACCTGAATCCTTTGATAGAGATAAGACAATAGCTTTAATCATGAATAGTTCAGGAAGCACAGGACTTCCTAAAGGTGTGGCACTTCCACATAGAACTGCTTGTGTTAGATTCTCTCATGCTAGAGATCCAATTTTTGGAAATCAAATAATTCCAGATACAGCAATACTAAGTGTAGTACCATTCCATCATGGATTTGGGATGTTTACAACTCTTGGATATTTAATTTGTGGTTTTAGAGTTGTATTAATGTATAGATTTGAGGAAGAACTCTTCCTTCGTTCACTACAAGACTATAAGATACAATCTGCTTTACTTGTACCAACTTTATTTTCATTTTTTGCTAAGAGTACTCTTATAGATAAATATGATTTAAGCAACCTGCATGAAATAGCATCAGGCGGCGCTCCACTATCTAAGGAAGTTGGAGAAGCTGTTGCTAAAAGATTCCACTTACCAGGAATCAGGCAGGGATATGGACTTACAGAAACAACTTCAGCAATTCTTATTACACCTGAAGGAGATGACAAGCCTGGAGCAGTAGGTAAAGTGGTACCATTCTTTGAAGCTAAAGT AGTAGATTTAGATACAGGAAAAACATTGGGAGTTAACCAGAGAGGAGAGCTGTGTGTAAGAGGACCTATGATAATGAGTGGATATGTAAATAATCCAGAAGCCACTAATGCATTAATAGATAAGGATGGATGGCTGCATTCTGGTGATATAGCATATTGGGATGAAGATGAACATTTTTTTATTGTAGATAGACTAAAATCCCTAATAAAATATAAGGGATACCAGGTAGCTCCAGCAGAATTAGAATCAATACTTCTGCAGCATCCAAACATATTTGATGCAGGAGTAGCTGGATTACCAGATGATGATGCAGGAGAACTTCCTGCTGCAGTAGTTGTTTTAGAGCATGGCAAAACTATGACTGAAAAAGAGATAGTTGACTATGTTGCAAGTCAGGTTACTACAGCAAAGAAATTGAGAGGCGGCGTAGTATTCGTAGATGAGGTTCCAAAAGGTCTTACAGGAAAATTGGATGCAAGAAAAATACGTGAAATACTTATAAAGGCAAAGAAGGGCGGCAAATCAAAATTATAA

##### *E. coli* codon-optimized

ATGGAAGACGCCAAAAACATAAAGAAAGGCCCGGCTCCATTCTATCCGCTAGAGGATGGAACCGCTGGAGAGCAACTGCATAAGGCTATGAAGAGATACGCCCTGGTTCCTGGAACAATTGCTTTTACAGATGCACATATCGAGGTGAACATCACGTACGCGGAATACTTCGAAATGTCCGTTCGGTTGGCAGAAGCTATGAAACGATATGGGCTGAATACAAATCACAGAATCGTCGTATGCAGTGAAAACTCTCTTCAATTCTTTATGCCGGTGTTGGGCGCGTTATTTATCGGAGTTGCAGTTGCGCCCGCGAACGACATTTATAATGAACGTGAATTGCTCAACAGTATGAACATTTCGCAGCCTACCGTAGTGTTTGTTTCCAAAAAGGGGTTGCAAAAAATTTTGAACGTGCAAAAAAAATTACCAATAATCCAGAAAATTATTATCATGGATTCTAAAACGGATTACCAGGGATTTCAGTCGATGTACACGTTCGTCACATCTCATCTACCTCCCGGTTTTAATGAATACGATTTTGTACCAGAGTCCTTTGATCGTGACAAAACAATTGCACTGATAATGAACTCCTCTGGATCTACTGGGTTACCTAAGGGTGTGGCCCTTCCGCATAGAACTGCCTGCGTCAGATTCTCGCATGCCAGAGATCCTATTTTTGGCAATCAAATCATTCCGGATACTGCGATTTTAAGTGTTGTTCCATTCCATCACGGTTTTGGAATGTTTACTACACTCGGATATTTGATATGTGGATTTCGAGTCGTCTTAATGTATAGATTTGAAGAAGAGCTGTTTTTACGATCCCTTCAGGATTACAAAATTCAAAGTGCGTTGCTAGTACCAACCCTATTTTCATTCTTCGCCAAAAGCACTCTGATTGACAAATACGATTTATCTAATTTACACGAAATTGCTTCTGGGGGCGCACCTCTTTCGAAAGAAGTCGGGGAAGCGGTTGCAAAACGCTTCCATCTTCCAGGGATACGACAAGGATATGGGCTCACTGAGACTACATCAGCTATTCTGATTACACCCGAGGGGGATGATAAACCGGGCGCGGTCGGTAAAGTTGTTCCATTTTTTGAAGCGAAGGTTGTGGATCTGGATACCGGGAAAACGCTGGGCGTTAATCAGAGAGGCGAATTATGTGTCAGAGGACCTATGATTATGTCCGGTTATGTAAACAATCCGGAAGCGACCAACGCCTTGATTGACAAGGATGGATGGCTACATTCTGGAGACATAGCTTACTGGGACGAAGACGAACACTTCTTCATAGTTGACCGCTTGAAGTCTTTAATTAAATACAAAGGATACCAGGTGGCCCCCGCTGAATTGGAGTCGATATTGTTACAACACCCCAACATCTTCGACGCGGGCGTGGCAGGTCTTCCCGACGATGACGCCGGTGAACTTCCCGCCGCCGTTGTTGTTTTGGAGCACGGAAAGACGATGACGGAAAAAGAGATCGTGGATTACGTCGCCAGTCAAGTAACAACCGCGAAAAAGTTGCGCGGAGGAGTTGTGTTTGTGGACGAAGTACCGAAAGGTCTTACCGGAAAACTCGACGCAAGAAAAATCAGAGAGATCCTCATAAAGGCCAAGAAGGGCGGAAAGTCCAAAT TGTAA

#### *C. autoethanogenum* metabolic genes

##### Acetolactate decarboxylase (CAETHG_RS14410)

MDDEVKVPNHIYQMSTINALVSGLYDGCVSLSKLLKKGNFGIGTFKGLDGELTLLNGTFYRTKPDGSVYVCSKNVSVPFAVVTELENYNTYNIQNRTSYEDIRKELDSFIESKNIFYAFYMEGKFNYVKTRTVVKQNMPYKPMAEVVKDQPMFEYNGVDGYVVGFRCPDYVEGLNVPGYHFHFINKDKKFGGHISEFSIENAKVYVQNCSCFRMELPKNESFYNMEVQDRNDEITSVEK*

##### Acetolactate synthase (CAETHG_RS08420)

MNRDIKKEVQLNTAQMLVKCLEAEGVKYIFGIPGEENLEIMNAISDSTIEFITTRHEQGAAFMADVYGRLTGKAGVCLSTLGPGATNLVTGVADADSDGAPVVAITGQVGTERMHITSHQFLDLCKMFEPITKRSKQIVRPDTVSEIIRLVFKYAESEKPGACHIDLPVNIAKMPVGALEKPLEKKIPPKEHADLSTIEEAASEIFKAKNPIILAGSGAIRGNSSKAVTEFATKLKIPVINTMMAKGIIPMDNKYSMWTIGIPQKDYVNKIIEEADLVITIGYDIVEYAPSKWNINGDIKIVHIDARPSHINKLYQPIVEVVGDISDALYNILRRTSSKDEPVKALEIKSEMLAEHESYANDNAFPMKPQRILNDVRKVMGPHDIVISDVGAHKMWIARHYNCYEPNTCIISNGFATMGIGVPGAIAAKLINPDKKVLAIVGDGGFMMNNQELETALRIKTPIVVLIFNDSNYGLIKWKQEEHYGKSCYVDFTNPDFVKLAESMYAKGYRVEKAEDLIPTLEEAFKQNVPAVIDCQVDYGENIKLTKHLKEVYENM*

##### Primary:secondary alcohol dehydrogenase (CAETHG_RS02620)

MKGFAMLGINKLGWIEKKNPVPGPYDAIVHPLAVSPCTSDIHTVFEGALGNRENMILGHEAVGEI AEVGSEVKDFKVGDRVIVPCTTPDWRSLEVQAGFQQHSNGMLAGWKFSNFKDGVFADYFHV NDADMNLAILPDEIPLESAVMMTDMMTTGFHGAELADIKMGSSVVVIGIGAVGLMGIAGSKLRG AGRIIGVGSRPVCVETAKFYGATDIVNYKNGDIVEQIMDLTHGKGVDRVIMAGGGAETLAQAVT MVKPGGVISNINYHGSGDTLPIPRVQWGCGMAHKTIRGGLCPGGRLRMEMLRDLVLYKRVDL SKLVTHVFDGAENIEKALLLMKNKPKDLIKSVVTF*

## References

Anastasina, M., Terenin, I., Butcher, S.J., Kainov, D.E., 2014. A technique to increase protein yield in a rabbit reticulocyte lysate translation system. Biotechniques 56. https://doi.org/10.2144/000114125

Brown, S.D., Nagaraju, S., Utturkar, S., De Tissera, S., Segovia, S., Mitchell, W., Land, M.L., Dassanayake, A., Köpke, M., 2014. Comparison of single-molecule sequencing and hybrid approaches for finishing the genome of Clostridium autoethanogenum and analysis of CRISPR systems in industrial relevant Clostridia. Biotechnol. Biofuels 7, 40. https://doi.org/10.1186/1754-6834-7-40

Bujara, M., Schümperli, M., Pellaux, R., Heinemann, M., Panke, S., 2011. Optimization of a blueprint for in vitro glycolysis by metabolic real-time analysis. Nat. Chem. Biol. 7, 271–277. https://doi.org/10.1038/nchembio.541

Carlson, E.D., Gan, R., Hodgman, C.E., Jewett, M.C., 2012. Cell-free protein synthesis: Applications come of age. Biotechnol. Adv. 30, 1185–1194. https://doi.org/10.1016/J.BIOTECHADV.2011.09.016

Caschera, F., Noireaux, V., 2014. Synthesis of 2.3 mg/ml of protein with an all Escherichia coli cell-free transcription–translation system. Biochimie 99, 162–168. https://doi.org/10.1016/J.BIOCHI.2013.11.025

Chappell, J., Jensen, K., Freemont, P.S., 2013. Validation of an entirely in vitro approach for rapid prototyping of DNA regulatory elements for synthetic biology. Nucleic Acids Res. 41, 3471–3481. https://doi.org/10.1093/nar/gkt052

Daniell, J., Nagaraju, S., Burton, F., Köpke, M., Simpson, S.D., 2015. Low-Carbon Fuel and Chemical Production by Anaerobic Gas Fermentation. Springer, Cham, pp. 293–321. https://doi.org/10.1007/10_2015_5005

Des Soye, B.J., Davidson, S.R., Weinstock, M.T., Gibson, D.G., Jewett, M.C., 2018. Establishing a High-Yielding Cell-Free Protein Synthesis Platform Derived from *Vibrio natriegens*. ACS Synth. Biol. 7, 2245–2255. https://doi.org/10.1021/acssynbio.8b00252

Des Soye, B.J., Gerbasi, V.R., Thomas, P.M., Kelleher, N.L., Jewett, M.C., 2019. A Highly Productive, One-Pot Cell-Free Protein Synthesis Platform Based on Genomically Recoded Escherichia coli. Cell Chem. Biol. 26, 1743–1754.e9. https://doi.org/10.1016/j.chembiol.2019.10.008

Dudley, Q.M., Nash, C.J., Jewett, M.C., 2019. Cell-free biosynthesis of limonene using enzyme-enriched Escherichia coli lysates. Synth. Biol. (Oxford, England) 4. https://doi.org/10.1093/SYNBIO/YSZ003

Endoh, T., Kanai, T., Imanaka, T., 2008. Effective approaches for the production of heterologous proteins using the Thermococcus kodakaraensis-based translation system. J. Biotechnol. 133, 177–182. https://doi.org/10.1016/J.JBIOTEC.2007.08.036

Endoh, T., Kanai, T., Imanaka, T., 2007. A highly productive system for cell-free protein synthesis using a lysate of the hyperthermophilic archaeon, Thermococcus kodakaraensis. Appl. Microbiol. Biotechnol. 74, 1153–1161. https://doi.org/10.1007/s00253-006-0753-3

Endoh, T., Kanai, T., Sato, Y.T., Liu, D. V., Yoshikawa, K., Atomi, H., Imanaka, T., 2006. Cell-free protein synthesis at high temperatures using the lysate of a hyperthermophile. J. Biotechnol. 126, 186–195. https://doi.org/10.1016/J.JBIOTEC.2006.04.010

Ezure, T., Suzuki, T., Shikata, M., Ito, M., Ando, E., 2010. A Cell-Free Protein Synthesis System from Insect Cells, in: Methods in Molecular Biology. Humana Press, pp. 31–42. https://doi.org/10.1007/978-1-60327-331-2_4

Failmezger, J., Scholz, S., Blombach, B., Siemann-Herzberg, M., 2018. Cell-Free Protein Synthesis From Fast-Growing Vibrio natriegens. Front. Microbiol. 9, 1146. https://doi.org/10.3389/fmicb.2018.01146

Feustel, L., Nakotte, S., Dürre, P., 2004. Characterization and Development of Two Reporter Gene Systems for Clostridium acetobutylicum. Appl. Environ. Microbiol. 70, 798–803. https://doi.org/10.1128/AEM.70.2.798-803.2004

Gan, R., Jewett, M.C., 2014. A combined cell-free transcription-translation system from *Saccharomyces cerevisiae* for rapid and robust protein synthe. Biotechnol. J. 9, 641–651. https://doi.org/10.1002/biot.201300545

Ghaffar, T., Irshad, M., Anwar, Z., Aqil, T., Zulifqar, Z., Tariq, A., Kamran, M., Ehsan, N., Mehmood, S., 2014. Recent trends in lactic acid biotechnology: A brief review on production to purification. J. Radiat. Res. Appl. Sci. 7, 222–229. https://doi.org/10.1016/J.JRRAS.2014.03.002

Gregorio, N.E., Levine, M.Z., Oza, J.P., 2019. A User’s Guide to Cell-Free Protein Synthesis. Methods Protoc. 2. https://doi.org/10.3390/mps2010024

Harbers, M., 2014. Wheat germ systems for cell-free protein expression. FEBS Lett. 588, 2762– 2773. https://doi.org/10.1016/j.febslet.2014.05.061

Heijstra, B.D., Kern, E., Koepke, M., Segovia, S., Liew, F., 2016. Novel Bacteria and Methods of Use Thereof. US 2016/0017276 A1.

Heijstra, B.D., Leang, C., Juminaga, A., 2017. Gas fermentation: cellular engineering possibilities and scale up. Microb. Cell Fact. 16, 60. https://doi.org/10.1186/s12934-017-0676-y

Hodgman, C.E., Jewett, M.C., 2013. Optimized extract preparation methods and reaction conditions for improved yeast cell-free protein synthesis. Biotechnol. Bioeng. 110, 2643– 2654. https://doi.org/10.1002/bit.24942

Hodgman, C.E., Jewett, M.C., 2012. Cell-free synthetic biology: Thinking outside the cell. Metab. Eng. 14, 261–269. https://doi.org/10.1016/J.YMBEN.2011.09.002

Jermutus, L., Ryabova, L.A., Plückthun, A., 1998. Recent advances in producing and selecting functional proteins by using cell-free translation. Curr. Opin. Biotechnol. 9, 534–548. https://doi.org/10.1016/S0958-1669(98)80042-6

Jewett, M.C., Calhoun, K.A., Voloshin, A., Wuu, J.J., Swartz, J.R., 2008. An integrated cell-free metabolic platform for protein production and synthetic biology. Mol. Syst. Biol. https://doi.org/10.1038/msb.2008.57

Jewett, M.C., Swartz, J.R., 2004. Mimicking theEscherichia coli cytoplasmic environment activates long-lived and efficient cell-free protein synthesis. Biotechnol. Bioeng. 86, 19–26. https://doi.org/10.1002/bit.20026

Jiang, Y., Liu, J., Jiang, W., Yang, Y., Yang, S., 2015. Current status and prospects of industrial bio-production of n-butanol in China. Biotechnol. Adv. 33, 1493–1501. https://doi.org/10.1016/J.BIOTECHADV.2014.10.007

Jones, D.T., 2005. Applied Acetone-Butanol Fermentation, in: Clostridia. Wiley-VCH Verlag GmbH, Weinheim, FRG, pp. 125–168. https://doi.org/10.1002/3527600108.ch5

Joseph, R.C., Kim, N.M., Sandoval, N.R., 2018. Recent Developments of the Synthetic Biology Toolkit for Clostridium. Front. Microbiol. 9, 154. https://doi.org/10.3389/fmicb.2018.00154

Karim, A.S., Dudley, Q.M., Juminaga, A., Yuan, Y., Crowe, S.A., Heggestad, J.T., Abdalla, T., Grubbe, W., Rasor, B., Coar, D., Torculas, M., Krein, M., Liew, F., Quattlebaum, A., Jensen, R.O., Stuart, J., Simpson, S.D., Köpke, M., Jewett, M.C., 2019a. In vitro prototyping and rapid optimization of biosynthetic enzymes for cellular design. bioRxiv. https://doi.org/10.1101/685768

Karim, A.S., Heggestad, J.T., Crowe, S.A., Jewett, M.C., 2018. Controlling cell-free metabolism through physiochemical perturbations. Metab. Eng. 45, 86–94. https://doi.org/10.1016/J.YMBEN.2017.11.005

Karim, A.S., Jewett, M.C., 2016. A cell-free framework for rapid biosynthetic pathway prototyping and enzyme discovery. Metab. Eng. 36, 116–126. https://doi.org/10.1016/J.YMBEN.2016.03.002

Karim, A.S., Rasor, B.J., Jewett, M.C., 2019b. Enhancing control of cell-free metabolism through pH modulation. Synth. Biol. ysz027. https://doi.org/10.1093/SYNBIO/YSZ027

Kay, J.E., Jewett, M.C., 2020. A cell-free system for production of 2,3-butanediol is robust to growth-toxic compounds. Metab. Eng. Commun. 10, e00114. https://doi.org/10.1016/j.mec.2019.e00114

Keasling, J.D., 2012. Synthetic biology and the development of tools for metabolic engineering. Metab. Eng. 14, 189–195. https://doi.org/10.1016/J.YMBEN.2012.01.004

Kelwick, R., Ricci, L., Mei Chee, S., Bell, D., Webb, A.J., Freemont, P.S., Freemont, P., 2017. Cell-free prototyping strategies for enhancing the sustainable production of polyhydroxyalkanoates bioplastics. bioRxiv. https://doi.org/10.1101/225144

Kelwick, R., Webb, A.J., MacDonald, J.T., Freemont, P.S., 2016. Development of a Bacillus subtilis cell-free transcription-translation system for prototyping regulatory elements. Metab. Eng. 38, 370–381. https://doi.org/10.1016/J.YMBEN.2016.09.008

Kim, D.M., Kigawa, T., Choi, C.Y., Yokoyama, S., 1996. A highly efficient cell-free, protein synthesis system from Escherichia coli. Eur. J. Biochem. 239, 881–886. https://doi.org/10.1111/j.1432-1033.1996.0881u.x

Kracke, F., Virdis, B., Bernhardt, P. V., Rabaey, K., Krömer, J.O., 2016. Redox dependent metabolic shift in Clostridium autoethanogenum by extracellular electron supply. Biotechnol. Biofuels 9, 249. https://doi.org/10.1186/s13068-016-0663-2

Kwon, Y.-C., Jewett, M.C., 2015. High-throughput preparation methods of crude extract for robust cell-free protein synthesis. Sci. Rep. 5, 8663. https://doi.org/10.1038/srep08663

Leuchtenberger, W., Huthmacher, K., Drauz, K., 2005. Biotechnological production of amino acids and derivatives: current status and prospects. Appl. Microbiol. Biotechnol. 69, 1–8. https://doi.org/10.1007/s00253-005-0155-y

Li, J., Wang, H., Jewett, M.C., 2018. Expanding the palette of Streptomyces-based cell-free protein synthesis systems with enhanced yields. Biochem. Eng. J. 130, 29–33. https://doi.org/10.1016/J.BEJ.2017.11.013

Li, J., Wang, H., Kwon, Y.-C., Jewett, M.C., 2017. Establishing a high yielding *streptomyces* - based cell-free protein synthesis system. Biotechnol. Bioeng. 114, 1343–1353. https://doi.org/10.1002/bit.26253

Liew, F., Henstra, A.M., Kӧpke, M., Winzer, K., Simpson, S.D., Minton, N.P., 2017. Metabolic engineering of Clostridium autoethanogenum for selective alcohol production. Metab. Eng. 40, 104–114. https://doi.org/10.1016/J.YMBEN.2017.01.007

Liew, F., Martin, M.E., Tappel, R.C., Heijstra, B.D., Mihalcea, C., Köpke, M., 2016. Gas Fermentation-A Flexible Platform for Commercial Scale Production of Low-Carbon-Fuels and Chemicals from Waste and Renewable Feedstocks. Front. Microbiol. | www.frontiersin.org 7. https://doi.org/10.3389/fmicb.2016.00694

Martin, R.W., Des Soye, B.J., Kwon, Y.-C.C., Kay, J., Davis, R.G., Thomas, P.M., Majewska, N.I., Chen, C.X., Marcum, R.D., Weiss, M.G., Stoddart, A.E., Amiram, M., Ranji Charna, A.K., Patel, J.R., Isaacs, F.J., Kelleher, N.L., Hong, S.H., Jewett, M.C.,. Cell-free protein synthesis from genomically recoded bacteria enables multisite incorporation of noncanonical amino acids_SI. Springer US.

Martin, R.W., Majewska, N.I., Chen, C.X., Albanetti, T.E., Jimenez, R.B.C., Schmelzer, A.E., Jewett, M.C., Roy, V., 2017. Development of a CHO-Based Cell-Free Platform for Synthesis of Active Monoclonal Antibodies. ACS Synth. Biol. 6, 1370–1379. https://doi.org/10.1021/acssynbio.7b00001

Meadows, A.L., Hawkins, K.M., Tsegaye, Y., Antipov, E., Kim, Y., Raetz, L., Dahl, R.H., Tai, A., Mahatdejkul-Meadows, T., Xu, L., Zhao, L., Dasika, M.S., Murarka, A., Lenihan, J., Eng, D., Leng, J.S., Liu, C.-L., Wenger, J.W., Jiang, H., Chao, L., Westfall, P., Lai, J., Ganesan, S., Jackson, P., Mans, R., Platt, D., Reeves, C.D., Saija, P.R., Wichmann, G., Holmes, V.F., Benjamin, K., Hill, P.W., Gardner, T.S., Tsong, A.E., 2016. Rewriting yeast central carbon metabolism for industrial isoprenoid production. Nat. Publ. Gr. 537. https://doi.org/10.1038/nature19769

Meinecke, B., Bertram, J., Gottsehalk, G., 1989. Purification and characterization of the pyruvate-ferredoxin oxidoreductase from Clostridium acetobutylicum, Arch Microbiol.

Mock, J., Zheng, Y., Mueller, A.P., Ly, S., Tran, L., Segovia, S., Nagaraju, S., Köpke, M., Dürre, P., Thauer, R.K., 2015. Energy Conservation Associated with Ethanol Formation from H2 and CO2 in Clostridium autoethanogenum Involving Electron Bifurcation. J. Bacteriol. 197, 2965–80. https://doi.org/10.1128/JB.00399-15

Moore, S.J., MacDonald, J.T., Wienecke, S., Ishwarbhai, A., Tsipa, A., Aw, R., Kylilis, N., Bell, D.J., McClymont, D.W., Jensen, K., Polizzi, K.M., Biedendieck, R., Freemont, P.S., 2018. Rapid acquisition and model-based analysis of cell-free transcription–translation reactions from nonmodel bacteria. Proc. Natl. Acad. Sci. 115, E4340–E4349. https://doi.org/10.1073/pnas.1715806115

Morgado, G., Gerngross, D., Roberts, T.M., Panke, S., 2018. Synthetic biology for cell-free biosynthesis: Fundamentals of designing novel in vitro multi-enzyme reaction networks, in: Advances in Biochemical Engineering/Biotechnology. Springer, Cham, pp. 117–146. https://doi.org/10.1007/10_2016_13

Nagaraju, S., Davies, N.K., Jeffrey, D., Walker, F., Köpke, M., Simpson, S.D., 2016. Genome editing of Clostridium autoethanogenum using CRISPR/Cas9. Biotechnol Biofuels 9, 219. https://doi.org/10.1186/s13068-016-0638-3

Nakamura, C.E., Whited, G.M., 2003. Metabolic engineering for the microbial production of 1,3-propanediol. Curr. Opin. Biotechnol. 14, 454–459. https://doi.org/10.1016/J.COPBIO.2003.08.005

Nakotte, S., Schaffer, S., Böhringer, M., Dürre, P., 1998. Electroporation of, plasmid isolation from and plasmid conservation in Clostridium acetobutylicum DSM 792. Appl. Microbiol. Biotechnol. 50, 564–567.

Nielsen, J., Fussenegger, M., Keasling, J., Lee, S.Y., Liao, J.C., Prather, K., Palsson, B., 2014. Engineering synergy in biotechnology. Nat. Chem. Biol. 10, 319–322. https://doi.org/10.1038/nchembio.1519

Nielsen, J., Keasling, J.D., 2016. Engineering Cellular Metabolism. Cell 164, 1185–1197. https://doi.org/10.1016/J.CELL.2016.02.004

Nirenberg, M.W., Matthaei, J.H., 1961. The dependence of cell-free protein synthesis in E. coli upon naturally occurring or synthetic polyribonucleotides. Proc. Natl. Acad. Sci. U. S. A. 47, 1588–602. https://doi.org/10.1073/pnas.47.10.1588

Rodriguez, B.A., Stowers, C.C., Pham, V., Cox, B.M., 2014. Green Chemistry PERSPECTIVE The production of propionic acid, propanol and propylene via sugar fermentation: an industrial perspective on the progress, technical challenges and future outlook. https://doi.org/10.1039/c3gc42000k

Salis, H.M., Mirsky, E.A., Voigt, C.A., 2009. Automated design of synthetic ribosome binding sites to control protein expression. Nat. Biotechnol. 27, 946–950. https://doi.org/10.1038/nbt.1568

Siegal-Gaskins, D., Tuza, Z.A., Kim, J., Noireaux, V., Murray, R.M., 2014. Gene Circuit Performance Characterization and Resource Usage in a Cell-Free “Breadboard.” ACS Synth. Biol. 3, 416–425. https://doi.org/10.1021/sb400203p

Silverman, A.D., Karim, A.S., Jewett, M.C., 2019a. Cell-free gene expression: an expanded repertoire of applications. Nat. Rev. Genet. https://doi.org/10.1038/s41576-019-0186-3

Silverman, A.D., Kelley-Loughnane, N., Lucks, J.B., Jewett, M.C., 2019b. Deconstructing Cell-Free Extract Preparation for *in Vitro* Activation of Transcriptional Genetic Circuitry. ACS Synth. Biol. 8, 403–414. https://doi.org/10.1021/acssynbio.8b00430

Swartz, J.R., Jewett, M.C., Woodrow, K.A., 2004. Cell-free protein synthesis with prokaryotic combined transcription-translation. Methods Mol. Biol. 267, 169–182. https://doi.org/10.1385/1-59259-774-2:169

Takahashi, Melissa K., Chappell, J., Hayes, C.A., Sun, Z.Z., Kim, J., Singhal, V., Spring, K.J., Al-Khabouri, S., Fall, C.P., Noireaux, V., Murray, R.M., Lucks, J.B., 2015. Rapidly Characterizing the Fast Dynamics of RNA Genetic Circuitry with Cell-Free Transcription– Translation (TX-TL) Systems. ACS Synth. Biol. 4, 503–515. https://doi.org/10.1021/sb400206c

Takahashi, Melissa K, Hayes, C.A., Chappell, J., Sun, Z.Z., Murray, R.M., Noireaux, V., Lucks, J.B., 2015. Characterizing and prototyping genetic networks with cell-free transcription-translation reactions. METHODS 86, 60–72. https://doi.org/10.1016/j.ymeth.2015.05.020

Tracy, B.P., Jones, S.W., Fast, A.G., Indurthi, D.C., Papoutsakis, E.T., 2012. Clostridia: the importance of their exceptional substrate and metabolite diversity for biofuel and biorefinery applications. Curr. Opin. Biotechnol. 23, 364–381. https://doi.org/10.1016/J.COPBIO.2011.10.008

Valgepea, K., de Souza Pinto Lemgruber, R., Meaghan, K., Palfreyman, R.W., Abdalla, T., Heijstra, B.D., Behrendorff, J.B., Tappel, R., Köpke, M., Simpson, S.D., Nielsen, L.K., Marcellin, E., 2017. Maintenance of ATP Homeostasis Triggers Metabolic Shifts in Gas-Fermenting Acetogens. Cell Syst. 4, 505–515.e5. https://doi.org/10.1016/j.cels.2017.04.008

Voloshin, A.M., Swartz, J.R., 2005. Efficient and scalable method for scaling up cell free protein synthesis in batch mode. Biotechnol. Bioeng. 91, 516–521. https://doi.org/10.1002/bit.20528

Wang, H., Li, J., Jewett, M.C., 2018. Development of a Pseudomonas putida cell-free protein synthesis platform for rapid screening of gene regulatory elements. Synth. Biol. 3. https://doi.org/10.1093/synbio/ysy003

Wang, S., Huang, H., Kahnt, J., Mueller, A.P., Köpke, M., Thauer, R.K., 2013. NADP-specific electron-bifurcating [FeFe]-hydrogenase in a functional complex with formate dehydrogenase in Clostridium autoethanogenum grown on CO. J. Bacteriol. 195, 4373–86. https://doi.org/10.1128/JB.00678-13

Wee, Y.-J., Kim, J.-N., Ryu, H.-W., 2006. Biotechnological Production of Lactic Acid and Its Recent Applications. Food Technol. Biotechnol. 44, 163–172.

Wiegand, D.J., Lee, H.H., Ostrov, N., Church, G.M., 2018. Establishing a Cell-Free *Vibrio natriegens* Expression System. ACS Synth. Biol. 7, 2475–2479. https://doi.org/10.1021/acssynbio.8b00222

Yim, H., Haselbeck, R., Niu, W., Pujol-Baxley, C., Burgard, A., Boldt, J., Khandurina, J., Trawick, J., Osterhout, R., Stephen, R., Estadilla, J., Teisan, S., Brett schreyer, H., Andrae, S., Hoon Yang, T., Yup Lee, S., Burk, M.J., Van dien, S., 2011. Metabolic engineering of Escherichia coli for direct production of 1,4-butanediol. Nat. Chem. Biol. 7. https://doi.org/10.1038/nCHeMBIO.580

Yim, S.S., Johns, N.I., Park, J., Gomes, A.L., McBee, R.M., Richardson, M., Ronda, C., Chen, S.P., Garenne, D., Noireaux, V., Wang, H.H., 2019. Multiplex transcriptional characterizations across diverse bacterial species using cell-free systems. Mol. Syst. Biol. 15, e8875. https://doi.org/10.15252/msb.20198875

